# Evolutionary Sequence and Structural Basis for the Epistatic Origins of Drug Resistance in HIV

**DOI:** 10.1101/2025.04.30.651576

**Authors:** Avik Biswas, Indrani Choudhuri, Kenneth Huang, Qinfang Sun, Andrej Sali, Ignacia Echeverria, Allan Haldane, Ronald M Levy, Dmitry Lyumkis

**Author notes:** A.B. and I.C. contributed equally to this work.

## Abstract

The emergence of drug resistance in the human immunodeficiency virus (HIV) remains a formidable challenge to the long-term efficacy of antiretroviral therapy (ART). A growing body of evidence highlights the critical role of epistasis, the dependence of mutational effects on the sequence context, in shaping the fitness landscape of HIV under ART-induced selection pressure. However, the biophysical origins of the epistatic interactions involved in engendering drug-resistance mutations (DRMs) remain unclear. Are the mutational correlations “intrinsic” to the properties of the protein, or do they arise because of drug binding? We use a Potts sequence-covariation statistical energy model built on patient-derived HIV-1 protein sequences to construct computational double mutant cycles that probe pairwise epistasis for all observed mutations across the three major HIV drug-target enzymes. We find that the strongest epistatic effects occur between mutations at residue positions that frequently mutate during the course of ART, termed resistance-associated positions. To investigate the structural origins of the strongest epistatic interactions, we perform ∼100 free energy perturbation molecular dynamics simulations, revealing that the primary contribution to the pairwise epistasis between DRMs arises from cooperative effects on protein stability and folding as an intrinsic consequence of the protein mutational landscape. The results collectively reinforce a mechanism of resistance evolution whereby viruses escape drug pressure by selectively engendering mutations at “intrinsically” coupled sites, allowing them to cooperatively ameliorate fitness detriments incurred by individual DRMs.

**Significance:** Epistasis refers to the phenomenon where the effect of a mutation on protein structure and function is dependent on the genetic sequence background of the mutation, resulting in the combined effect of mutations being non-additive. Epistasis plays a significant role in the evolution of drug resistance in viruses such as HIV under therapeutic selection pressure. We combine a protein sequence coevolutionary model and molecular dynamics free energy simulations to identify and probe the mechanistic origins of the strongest epistatic interactions connecting HIV drug-resistance mutations. The work establishes a foundation to probe the molecular bases of epistasis and predict the evolution of resistance predicated on the knowledge of epistatic interaction networks.

## Introduction

The rapid emergence of drug resistance in the human immunodeficiency virus (HIV) remains a formidable challenge to the long-term efficacy of antiretroviral therapy (ART) [1-3]. A growing body of evidence highlights the critical role of epistasis, the dependence of mutational effects on the sequence context, in modulating the fitness landscapes of viruses such as HIV under selection pressure [4-7]. As a consequence of epistasis, the effect of a single mutation on protein structure and function is contingent on the genetic sequence background of the mutation that includes residue(s)/mutation(s) at other proximal and/or distal sites. The result is that the combined effect of mutations is often non-additive [8, 9].

Epistasis leads to correlated mutations in viral proteins observed under external ART selection pressure, which offers viable escape pathways while maintaining replication competent viruses [10-12]. In the context of HIV drug resistance, correlated mutations have been extensively studied and tabulated in large public databases, such as the Stanford HIV drug resistance database (HIVDB) [13-15]. However, the biophysical origins for how mutational correlations lead to functional adaptations such as drug resistance are not known. Mutations can interact because of physical, structural, or functional couplings at the molecular level. Evolutionary adaptation can rewire intramolecular interactions via epistasis highlighting that mutation can reconfigure structural interaction networks, leading to non-additive functional effects [16]. The central question we aim to answer is whether the mutational couplings observed in HIV under ART selection are an intrinsic property of the protein’s mutational landscape, or if they emerge as a direct result of cooperative effects induced by drug binding. We refer to the inherent structural or functional interactions that originate in exclusion of any external perturbation, even in case where such perturbations result from drug selection pressure as “intrinsic”. In contrast, correlations that originate as a direct consequence of external perturbations, particularly due to cooperative effects induced by drug binding, [17] [18, 19], provide an alternative path to drug resistance.

The pairwise epistasis between mutations can be studied using double mutant cycles (DMCs) constructed to quantify the non-additivity of fitness effects between double and individual single mutations. Introduced first in context of protein engineering [17], DMCs probe the strength of epistatic interactions by comparing the effects of single and double mutations on specific molecular properties, such as binding affinity, enzymatic activity, or structural stability. To obtain the DMC effects on HIV evolutionary fitness, we use a Potts statistical co-evolutionary model based on patient-derived HIV protein sequences from the Stanford HIVDB. Potts models have demonstrated accuracy in predicting viral fitness landscapes [20-24], including HIV evolution under ART selection pressure [4-7, 18, 19, 25]. By capturing the co-evolutionary signals, the Potts model provides a prevalence-based measure of evolutionary fitness and allows for efficiently scanning large numbers of DMCs computationally [25], Conventional experimental matrices of epistasis remain intractable, due to the combinatorically vast space of potential mutation pairs and the resource-intensive phenotypic assays required for validation. Sequence-based Potts models also have significant advantages over other measures of epistasis. Current strategies to identify correlated mutations emerging under ART rely predominantly on clinical observations from patient datasets. However, the multifaceted nature of correlated interactions makes it difficult to distinguish between coincidental co-occurrence and true cooperativity [26]. Thus, computational approaches emerge as indispensable tools, to systematically identify and interrogate epistatic networks which underpin primary and secondary mutations responsible for HIV drug resistance. The vast repositories of patient-derived HIV sequence data available in large-scale public databases also offers important opportunities, as Potts models derived from drug-experienced patient sequences can effectively identify and elucidate the epistatic interaction networks underlying evolving HIV drug resistance. However, the Potts model is unable to distinguish between the types of cooperativities which lead to drug resistance under ART treatment.

A major source of epistasis in proteins is the constraint to maintaining a valid, functional fold. *Posfai et al.* [27] have demonstrated that selection for protein thermodynamic stability selects for epistatic interactions that are necessary to drive folding [27, 28]. To probe whether the mutational correlations under ART selection relate to intrinsic effects on protein stability, we use structure-based free energy perturbation molecular dynamics (FEP/MD) to quantify the thermodynamic effects of mutations on protein stability. Recent advancements in FEP/MD methodologies have demonstrated their ability to surpass empirical scoring functions and machine learning models in predicting mutational effects [29], particularly in complex, highly dynamic systems, like viral proteins under drug selection pressure [30, 31]. Recent work [32-34] has also demonstrated that FEP/MD cycles can be specifically designed to capture the epistatic effects of protein mutations, probe the molecular mechanisms underlying epistatic interactions, and guiding rational drug design. We present a framework rooted in protein biophysics that combines sequence-based Potts co-evolutionary models with structure-based FEP molecular dynamics to probe the evolutionary sequence and structural basis for the epistatic origins of drug resistance in HIV.

Deciphering the biophysical basis of epistasis is crucial to linking protein sequence, structure, and function, and in the context of HIV, elucidating the mechanistic underpinnings of drug resistance and guiding therapeutic design strategies. Using HIV as a model system, we reveal that the primary contribution to the strongest epistatic couplings driving the evolution of drug resistance arise from intrinsic effects on protein stability, supporting a mechanism of resistance evolution through correlated mutations at intrinsically coupled residue sites that offer viable escape pathways while maintaining or ameliorating potentially harmful effects on structural integrity [4].

## Results

Owing to its high mutation rate [35, 36], rapid replication cycle [36, 37], strong selective pressure of ART therapies [36], and extensive data availability in public repositories such as the Stanford HIVDB, HIV provides a rich model system with which to study drug resistance dynamics and evolution. We focus on the three viral enzymes, protease (PR), reverse transcriptase (RT), and integrase (IN). PR functions as a homodimer of two p12 subunits to cleave viral gag and gag-pol polyproteins during viral maturation. RT functions as a heterodimer of p51 and p66 subunits and converts the viral RNA into complementary viral DNA to prepare the viral genome for chromatin integration following ingress. IN functions within a large nucleoprotein assembly called an intasome and mediates the catalytic integration of viral DNA into host chromatin to establish infection in target cells. All three HIV enzymes are also targets of ART [38]. PR function is blocked by PIs [5], which engage the catalytic site of the PR dimer and inhibit enzymatic activity. RT function can be blocked either by the nucleoside reverse transcriptase inhibitors (NRTIs), which directly cause chain termination during DNA synthesis, or by the non-nucleoside reverse transcriptase inhibitors (NNRTIs), which inhibit DNA synthesis through allosteric effects. IN function is blocked by the integrase strand transfer inhibitors (INSTIs), which inhibit the incorporation of viral DNA into host target DNA.

### Epistasis in HIV evolution under drug selection pressure

Experimental techniques to assess the effect of multiple mutations on phenotype have proven effective [39-41], but functional assays to test all possible combinations remain inaccessible. Therefore, robust computational methods to predict the combinatorial effects of mutational variations are needed to properly infer epistasis. For HIV, large patient-derived sequence datasets (Stanford HIVDB) can be used to derive co-evolutionary information to serve as a basis for building Potts Hamiltonian models of protein structure and function [20, 21, 42-45]. The Potts model is a maximum-entropy model inferred from a multiple sequence alignment (MSA) of related protein sequences, constrained to capture the bivariate (pairwise) residue marginals between all pairs of positions in the MSA. In the Potts Hamiltonian formalism, a central quantity known as the ‘statistical’ energy of a sequence *S* (see Methods) is interpreted to be proportional to evolutionary fitness; the model predicts that sequences will appear in the dataset with probability, *P*(*S*) ∝ *e*^−*E*(*S*)^, such that favorable statistical energies are indicative of higher prevalence. The Potts model has been successfully used to predict the tertiary and quaternary structures of proteins [42, 46-53] as well as viral protein stability and fitness landscapes [5-7, 19, 20, 22, 23, 45, 54-56].

We use the Potts model as a scanning tool to calculate DMCs between every pair of mutations in the three primary drug target proteins – PR, RT, and IN. The Potts model provides a log likelihood for any mutation *M*_*i*_ at a position *i* in a given sequence background relative to the wild-type residue (consensus), which can be quantified by the resulting change in the Potts statistical energy due to the mutation, 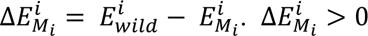 implies a favorable mutation likely accompanied by a fitness penalty for reversion (in the given background), and *vice versa*. The DMC for a pair of mutations, *M*_*i*_, *M*_*j*_ can be estimated as the difference in log-likelihoods (Potts Δ*E*) between the double mutant and the sum of the effects for the individual single mutants as 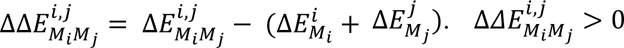 implies that the pair of mutations are positively or synergistically epistatically coupled in fitness effects; conversely, 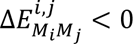 implies negative epistasis or antagonistically coupled mutations. 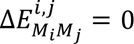 implies zero epistasis, which is equivalent to simple additivity.

**Fig. 1** illustrates the distribution of DMC effects (ΔΔ*E*) in HIV-1 PR, RT, and IN, respectively. A vast majority of mutational effects are clustered around 0, implying additivity in fitness effects, and only a small fraction of the mutation pairs exhibits strong epistasis (the tail ends of the distribution in **Fig 1**). However, within the small number of mutation pairs exhibiting the strongest signatures of pairwise epistasis, many of them involve mutations at drug resistance associated positions (DRAPs). The majority (>>50%) of the 50 strongest DMCs in PR and RT involve mutations at DRAPs, whereas in IN, ∼40% of the 50 strongest DMCs involve mutations at DRAPs (**Supplementary Information, Section S1 and Table S1**). These results are indicative of the role of epistasis in giving rise to correlated mutations under drug selection pressure eventually leading to the evolution of resistance. *Butler et al.* [4] have shown that knowledge of the strongest coupled positions inferred using a drug-naive Potts model based on patient-derived HIV protease sequences prior to ART initiation can be used to capture the strongest epistatically coupled sites that successfully identify the residue positions where DRMs arise in HIV PR. However, a drug-naïve Potts model, even if successful in predicting DRAPs, cannot be used to capture the epistatic interactions connecting DRMs. This is because DRMs appear at very low (or zero) frequencies in drug-naïve datasets, and provide insufficient statistics to capture the epistatic effects (Methods).

**Fig. 1:**
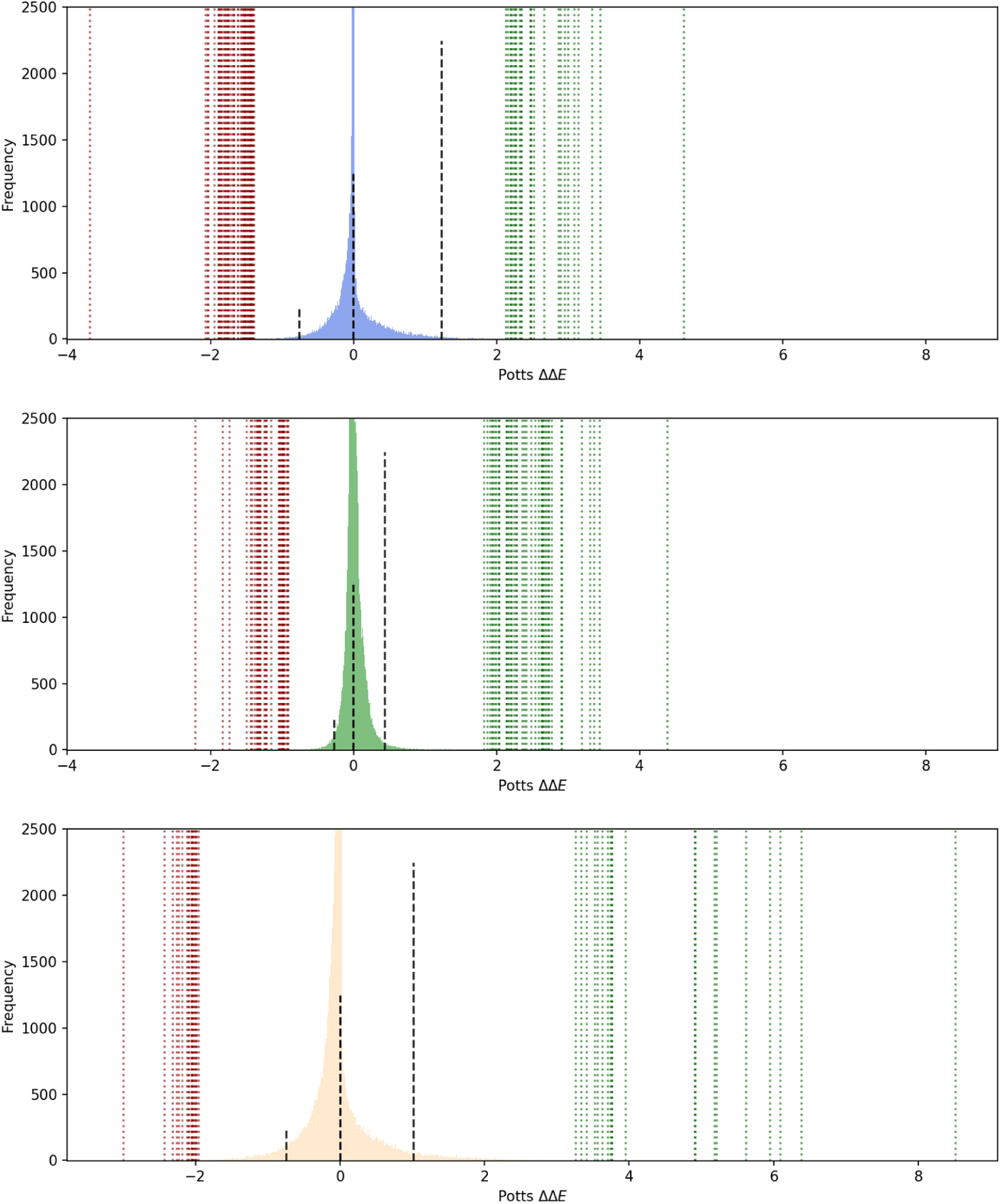
Potts model scan of double mutant cycles. Figure shows the Potts model scan of all possible DMCs in HIV-1 PR (blue) in top panel, RT (green) in middle, and IN (wheat) in the bottom panel. Black dashed lines indicate the 1^st^, 50^th^, and 99^th^, quartiles, respectively. Figures are shown only to an ordinate limit of 2500, from more than 100,000 indicating the vast majority of mutational interactions are close to zero(0), or non-epistatic. Red, and green dotted vertical lines indicate the strongest double mutant cycles, antagonistic, and synergistic, respectively at RAPs from within the top 50 cycles in each case.

When the environment in which HIV replicates changes due to ART application, HIV must mutate to abrogate drug binding while simultaneously preserving protein function. The presence of large epistatic interactions within all three viral enzymes implies that viral proteins are able to co-mutate to engender evolutionary pathways to ameliorate the substantial fitness cost of individual DRMs. However, the fitness penalty can be compensated through viable secondary mutations that cooperatively couple with the DRM, thereby inducing drug resistance without significant detriment to the virus. These synergistically coupled sites are therefore more likely to be associated with resistance. Here, our assumption is that resistance cannot be achieved through selectively neutral mutations at single sites, as drug treatment would likely be ineffective. This indeed appears to be the case for HIV protease, where single DRMs are usually deleterious [57]. We accordingly suggest that this principle applies to other enzymatic drug targets, including RT and IN.

### Potts model double mutant cycles reveal the strongest epistatic interactions affecting drug resistance

In this section, we identify the strongest DMCs between pairs of mutations that involve at least one DRAP in each of the three predominant drug-target proteins in HIV. **Table 1A** shows the strongest synergistic (ΔΔ*E >* 0) pairs of double mutations where at least one amongst the pair is classified as a “primary” DRM (see Methods). Synergistic effects can encompass compensatory effects on viral fitness, protein stability, enzymatic activity, as well as phenotypic effects such as increased resistance profiles [14].

**Table 1:**
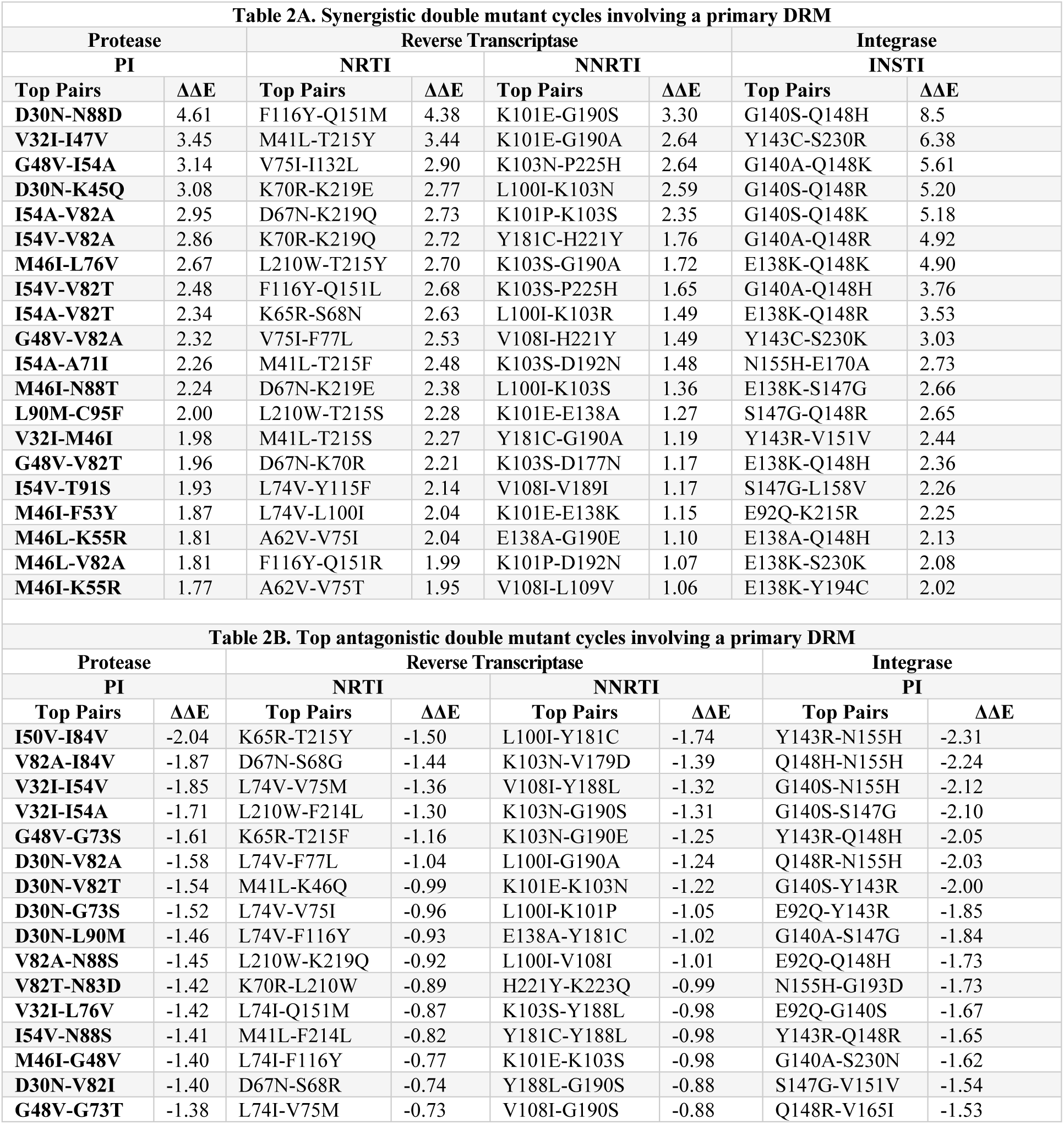
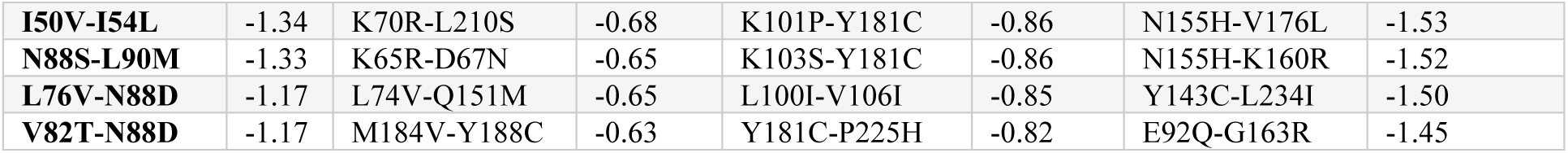
(**A**). Potts model predictions of the top 20 synergistic double mutant cycles where the pair first mutation of the pair is categorized as a “primary” DRM appearing with >1% frequency. The table shows the top pairs of mutations where the DMC effect is predicted to be most synergistic (ΔΔE > 0). The second mutation in the pair can be any other mutation in the same protein. Many of the predicted top pairs are both clinically observed resistance mutation pairs known to occur together with synergistic effects. (**B**). Potts model predictions of the top 20 antagonistic DMCs where the first of the pair is categorized as a “primary” DRM appearing with >1% frequency. The second mutation in the pair can be any other mutation in the same protein as the first mutation.

Many of the synergistic DMC pairs that we identify recapitulate characterizations of compensatory behavior in the literature (**Supplementary Information, Section S2**). In addition to identifying known DRM combinations for each of the three drug targets, our computational models are also able to distinguish between distinct evolutionary pathways within a given protein. For example, there are two distinct types of thymidine analogue mutations (TAMs) that affect the susceptibility of RT to NRTIs. Our findings recapitulate the known DRM pairs selected by NRTIs, and also include a previously unknown pair (L74V-L100I) amongst the strongest DMCs. L74V is an NRTI-associated DRM, whereas L100I is an NNRTI-associated DRM. Presence of both mutations in response to different drug classes suggests a high likelihood of pan-drug resistance. Furthermore, co-occurring DRM combinations from the Stanford HIVDB were also identified as being strongly synergistic DMCs in our analyses. Of the reported co-occurring mutations pairs in the Stanford HIVDB, a number (PR: ∼88%, RT: ∼78%, IN: ∼90%) of mutations pairs present were found to provide synergistic epistatic interactions (positive DMC values). This further supports the hypothesis that evolution of drug resistance capitalizes on pre-existing coupled interactions in a manner that has a highly non-linear effect on protein fitness [4].

The Potts model can also be utilized to probe DMCs between mutation pairs involving DRMs and those exhibiting strong antagonistic behavior (**Table 1B**). Unlike synergistic pairs, antagonistic pairs interfere with one another, thereby co-occur infrequently and accordingly are not well represented in the literature. However, we found empirical evidence of antagonism that agrees with the DMCs identified through our Potts model. For example, treatment with the two NRTIs (3TC, AZT) are known to decrease cross-resistance [58, 59], with treatments combining both showing higher sustained efficacies than treatment with a single inhibitor [60, 61]. In agreement, we find very strong antagonism between K65R, which confers resistance to 3TC, and T215Y, which arises with treatment of AZT. K65R is also known to exhibit bidirectional antagonism with TAMs [62-65]. As discussed below, inhibitors targeting sites with residues exhibiting such behavior can open novel design space for future in the face of rising drug resistance, either when given in combination or sequentially to retain drug susceptibility.

### Therapeutic recommendations and design strategies for combinatorial ART

In this section, we identify strongly coupled pairs using the full-range DMC scan with the Potts sequence-covariation model, providing a template to guide combinatorial ART (c-ART) to avoid/exploit pre-existing couplings in the evolutionary landscape. Only pairs comprised of individual primary DRMs were considered, with the selection criteria being that (i) the DRMs must respond to distinct chemical subspaces in different drugs but not both, and (ii) pairs must be strongly coupled (*P_1:50_*) in synergistic or antagonistic epistasis identified via the Potts model DMCs (Methods). This selection criteria enables the identification of DRMs that can be pragmatically avoided, or targeted for effective c-ART. Therefore, pairs involving two DRMs which affect different drugs within the same class, multiple classes of drugs, or have overlap in selection by multiple drugs, were avoided in this analysis.

Application of c-ART drugs which have correspondent DRMs that are synergistically coupled in the Potts model should be avoided, as the compensatory effect can lower the fitness barrier to accelerate resistance development. Single DRMs bias the fitness landscape to predispose certain stabilizing secondary mutations; addition of another drug exerts selective pressure which expedites the occurrence of the secondary DRM. **Table 2A** shows strongly synergistic DRM combinations that are co-selected by different drugs. We largely focus on the NRTI and NNRTI classes of RT inhibitors, which have been historically used in combination, and chronological sequence [66]. We have also listed specific drugs that might elicit unwarranted synergistic DRMs in each case. These synergistic couplings between DRMs selected by these classes (NRTIs and NNRTIs) were previously unknown. Importantly, we recapitulate an important example of a previously known synergistic effect (Q151M-F116Y) in RT resulted from treatment of NRTIs [62]. In PR, a novel synergistic pair (V32I-I47V) can also co-occur over the course of treatment and likely elicit a similar response of lowering the barrier to PI drug resistance.

**Table 2A.**
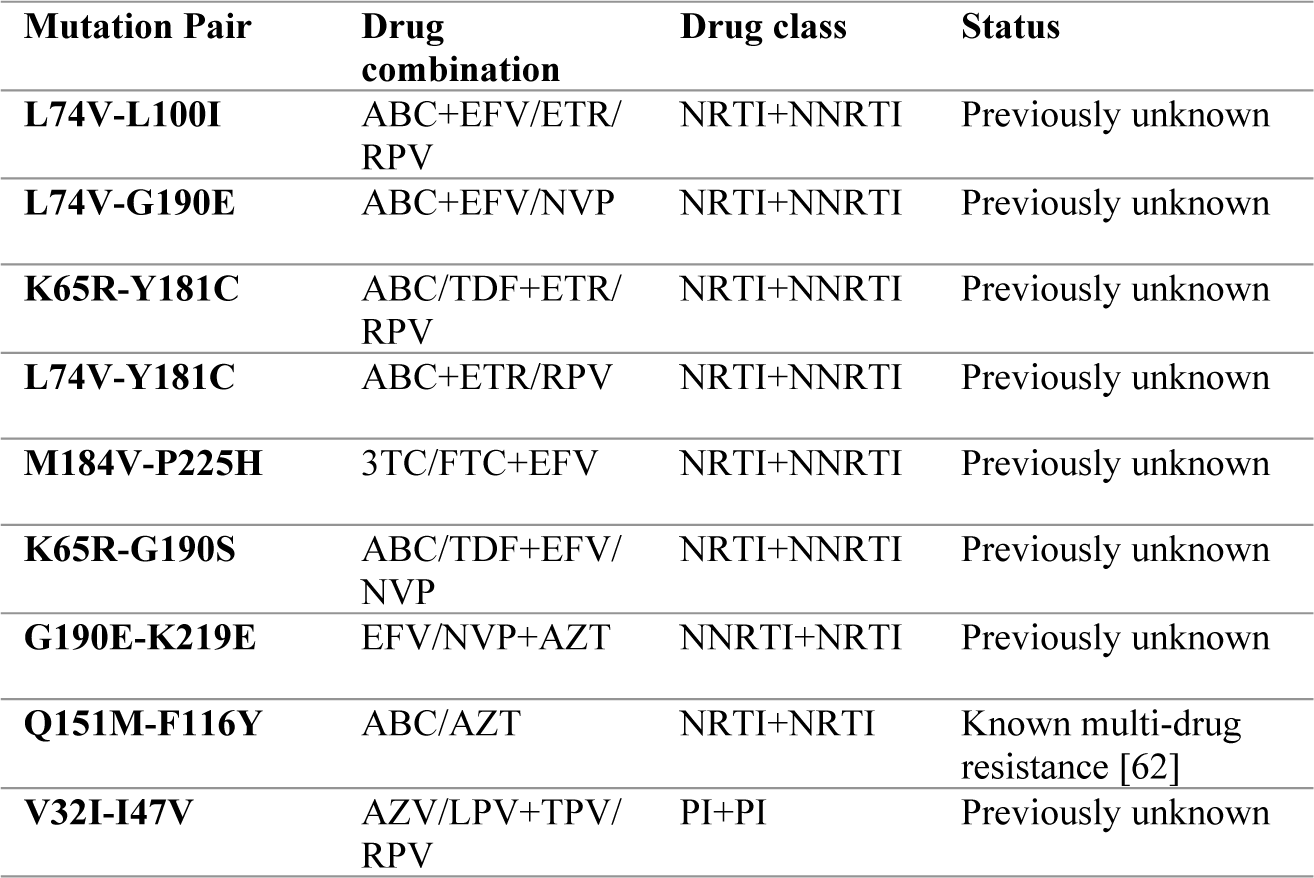
Mutation Combinations to Avoid due to synergistic couplings.

**Table 2B.**
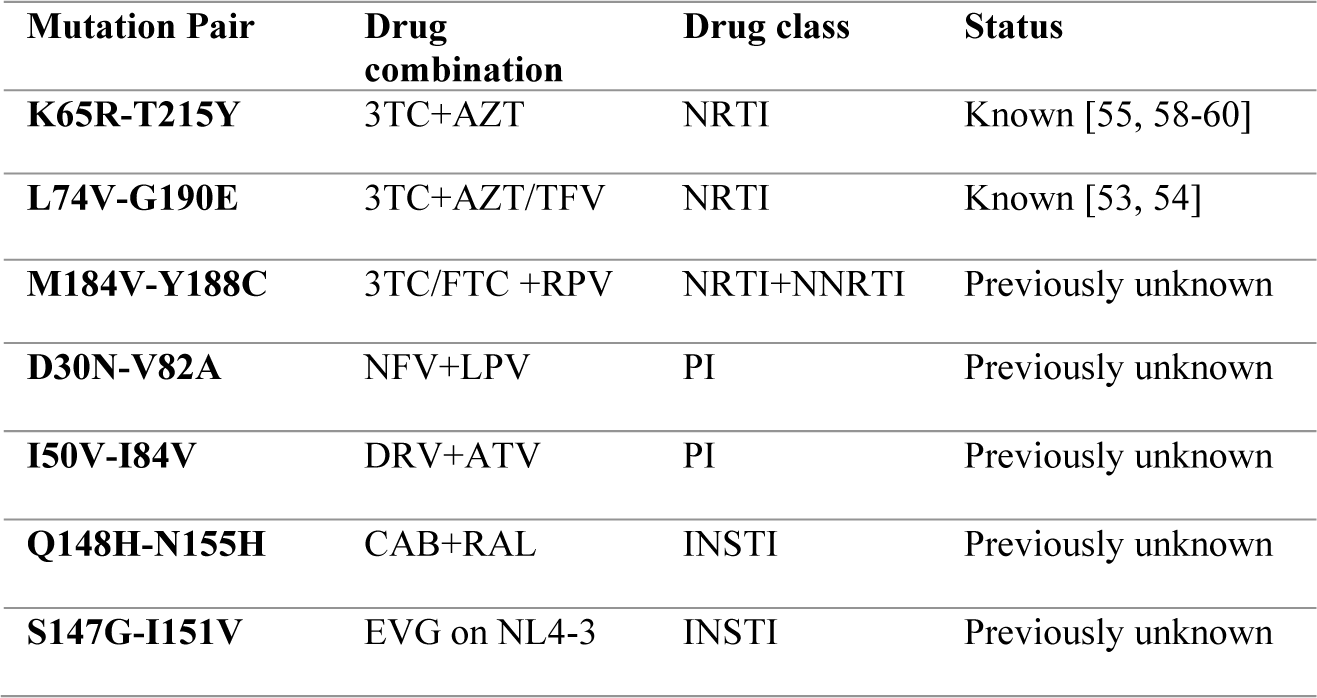
Mutation Combinations to Target due to antagonistic couplings.

| **Mutation Pair** | **Drug combination** | **Drug class** | **Status** |
| --- | --- | --- | --- |
| **K65R-T215Y** | 3TC+AZT | NRTI | Known [55, 58-60] |
| **L74V-G190E** | 3TC+AZT/TFV | NRTI | Known [53, 54] |
| **M184V-Y188C** | 3TC/FTC +RPV | NRTI+NNRTI | Previously unknown |
| **D30N-V82A** | NFV+LPV | PI | Previously unknown |
| **I50V-I84V** | DRV+ATV | PI | Previously unknown |
| **Q148H-N155H** | CAB+RAL | INSTI | Previously unknown |
| **S147G-I151V** | EVG on NL4-3 | INSTI | Previously unknown |

Prior work has indicated that DRMs in HIV can become evolutionarily “entrenched” during the course of ART treatment, substantially increasing reversion penalty to the parent wild-type even after therapy withdrawal [5, 6]. This entrenchment has been shown to play a central role in the persistence of drug-resistant viruses, and many strongly entrenched mutations have been shown to revert slowly in patients after ending ART [5, 67-69]. Therefore, past treatment regimens can result in persistent populations with DRMs in significant representation, increasing the overall likelihood of a secondary DRM occurring in presence of a secondary ART treatment, even when applied at a later time point. Many of the DRMs selected by multiple ART classes, with exception of NNRTIs, are highly entrenched in the patient population [6]. We thus propose that drugs selecting for mutations coupled synergistically should also be avoided in chronological or sequential order, as to avoid sub-optimal outcomes.

It is also possible that specific drug combinations can lead to a genetically defined virus with altered fitness and resistance profiles such that the virus remains susceptible to drugs in the current or next ART regimen. One avenue through which an adaptation such that a virus resistant to one drug will exhibit a lower barrier to resistance to a different drug, can be achieved is by using ART drugs selecting for DRMs that exhibit strong antagonistic couplings. Our general hypothesis is that inhibitors targeting residue sites exhibiting such antagonistic phenotypes can open exciting possibilities of hindering the development of drug resistance both when used in combination as part of c-ART or in chronological sequence order, such that virus resistant to one drug remains susceptible to the other drug.

**Table 2B** shows the strong antagonistic couplings that can be exploited for c-ART. In support of our hypothesis, we note that the NRTIs, 3TC and AZT are known to select for K65R-T215Y and L74V-G190E, which are antagonistically coupled mutation pairs. When both pairs of mutations arise, resistance to one drug decreases resistance to the other [58, 59], leading to higher sustained efficacies using AZT+3TC combinations [60, 61]. We uncover previously unknown antagonistically coupled DRM pairs that can be targeted by distinct drugs in combination or sequential order (**Supplementary Information, Section S3, Figure S1**). We hypothesize that these antagonisms can be utilized for drug design strategies. Drugs can be designed to utilize interactions with residues positions that are antagonistically coupled while avoiding residue positions of known synergistic couplings such that mutations induced at one site cannot be compensated for by mutations at another interacting site.

### Molecular Dynamics simulations reveal the biophysical basis of strongest epistatic interactions modulating drug resistance

The Potts statistical energy model provides a sequence-based framework for analyzing the cooperative effects of mutations on fitness, and it can be used to rapidly scan for the strongest mutational correlations. We now use a structure-based framework, alchemical free energy perturbation molecular dynamics (FEP/MD) simulations, to quantify the thermodynamic effects on protein stability of those mutation pairs identified by the Potts model as the most highly cooperative. By simulating the atomic-level changes associated with mutating protein residues, FEP/MD provides accurate estimates of the free energy differences that capture the structural and energetic perturbations induced by mutations in the protein system [70, 71]). **Fig 2A** depicts an alchemical free energy thermodynamic cycle used to determine the difference in the free energy change (ΔΔG) of protein unfolding due to mutation from the wild-type (WT) to a mutant (M) protein state, involving both folded and unfolded states for each of the WT and mutant forms. The free energy difference, ΔG(WT), represents the transition from the folded to the unfolded state in the WT protein, while ΔG(M) represents the same transition in the mutant protein. By completing this cycle, the difference in the free energy change (ΔΔG) associated with unfolding due to the mutation can be obtained by mutating the WT residue to the mutant in both the folded and unfolded states. This “alchemical” free energy path can be used quantify the stabilizing or destabilizing impact of the mutation on protein folding[70, 71].

**Fig 2:**
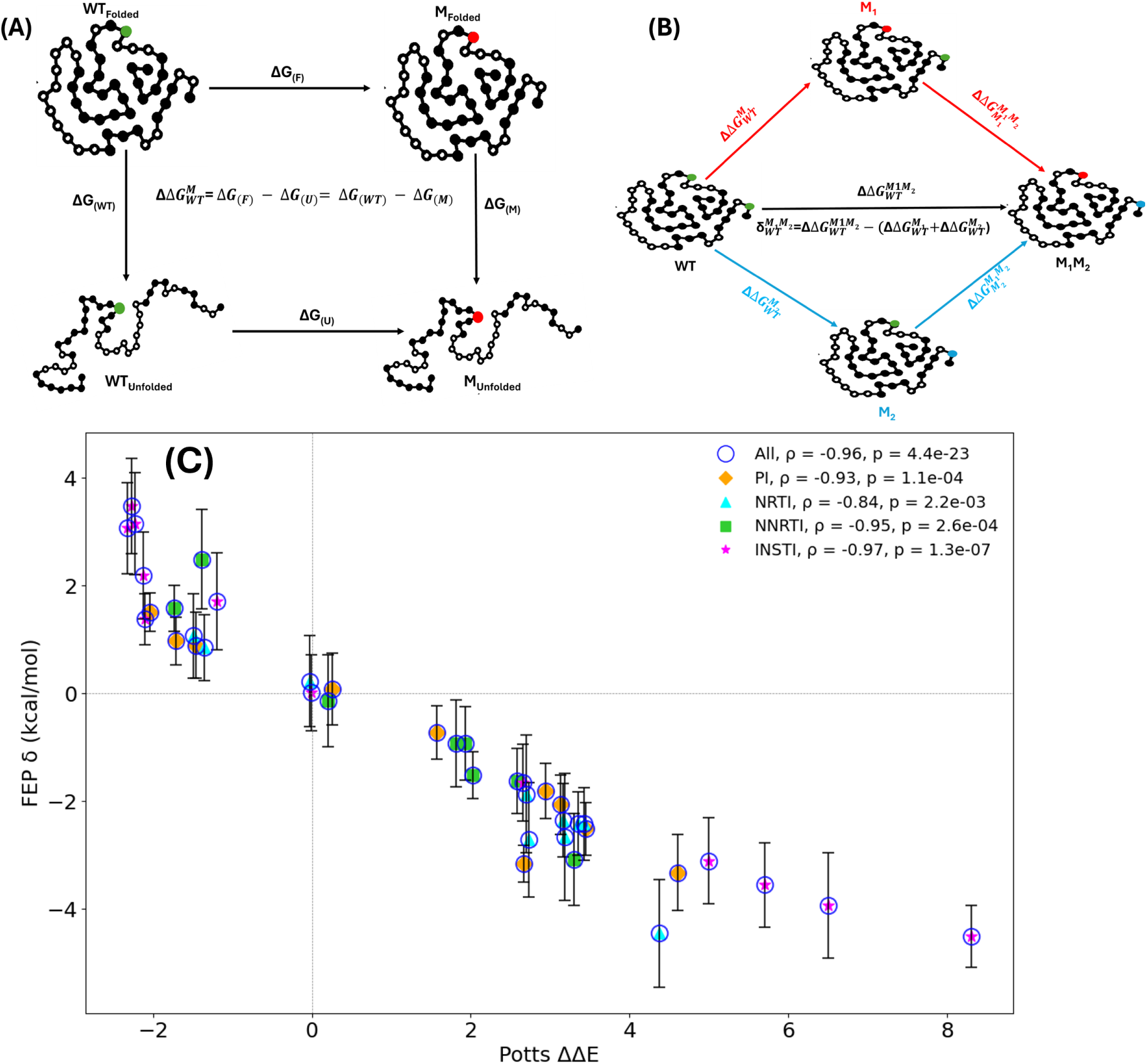
Thermodynamic cycles of Mutational free energy calculations and correlation between structure-based FEP/MD and evolutionary sequence-based Potts model. (A) Alchemical free energy calculation cycle for the calculation of the ΔΔG of unfolding upon amino acid mutation. ΔG(M) and ΔG(WT) denote the free energy changes. (B) Double mutant cycle with three different pathways to calculate a double mutation free energy and the corresponding non-additivity. The subscripts in the equation denote the reference state, while superscripts show the mutational target state. (C) Figure shows the correlation between structure-based FEP/MD double mutant cycles (δ) and the Potts model predictions (ΔΔE) for the strongest correlated mutation pairs such that one amongst the mutation is a primary DRM. 40 mutation pairs across PR, RT, and IN, are shown as blue circles and error bars are indicated in black. Individual mutation pairs from the 3 proteins selected by the 4 different classes of drugs, PIs, NRTIs, NNRTIs, and INSTIs are indicated with individual markers, and the corresponding Spearman rank-order correlation coefficients are shown in the legend. Correlation is > |0.90| for all except for the NRTI class of drugs (correlation =-0.84), with a p-value < 0.001 indicating statistical significance.

For double mutants, an additional thermodynamic cycle, illustrated in **Fig 2B** incorporates both single and double mutant states designed along the lines of [32, 34] to investigate the non-additive effects associated with combined mutations. The approach, developed by *Werner et al.* [32], demonstrated that alchemical free energy molecular dynamics simulations can accurately reproduce cooperative thermodynamic changes in protein stability associated with correlated pairs of mutations. The double mutant cycle (DMC) constructed in **Fig 2B** enables us to isolate and quantify the deviation from additivity (δ), which reflects the epistatic interactions between mutations. Structurally, such non-additivities arise from direct side-chain interactions, allosteric communication, or conformational rearrangements that can affect both local and/or distal regions of the protein. In the context of HIV enzymes, applying DMC allows us to determine how specific combinations of DRMs, even when distantly located, might cooperatively reshape the protein’s energetic landscape. This not only provides a molecular rationale for the epistatic coupling but also helps uncover synergistic or antagonistic mutations that modulate intrinsic fitness, offering structural insights that go beyond sequence-level correlations.

Because free energy is a state function, the cycle in **Fig 2B** provides three equivalent pathways for computing the double mutation free energies offering flexibility in calculating cooperativity between residues. Substituting different pathways into the equation in **Fig 2B** provides three possible estimates for non-additivity 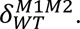 Notably, contributions from the unfolded state are canceled out due to the assumption of additivity in substitutions in the unfolded protein. As a result, non-additivities are determined solely from the folded state. There are 3 different paths to achieve double mutations from WT indicated with red, black and blue color in Fig. 2B. We use a single pathway (black, middle, **Fig. 2B**) that directly connects the WT to the double mutant M1M2, along with the free energy changes associated with the single mutant transformations from WT to M1 (red, top, Fig. 2B) and WT to M2 (blue, bottom, Fig. 2B), to estimate the non-additivities

To investigate the mutational effects on protein stability, we choose 40 of the top coupled mutation pairs that involve a primary DRM appearing at >1% frequency in drug-experienced patient samples in the Stanford HIVDB. We used apo structures to initiate the alchemical FEP molecular dynamics simulations of the mutations; we did <u>not</u> perform FEP simulations of the mutational effects on ligand binding affinities. We observe very strong correlation (with high statistical significance) between the sequence-based Potts model predictions of epistatic effects on protein fitness and the cooperative effects on protein stability obtained using structure-based FEP/MD. **Fig 2C** illustrates the correlation between all 40 mutation pairs across the 3 target proteins in *Pol*, as well as individually across each protein for 4 different drug classes.

The strong correlation in **Fig 2C** establishes the effectiveness of using evolutionary sequence data to predict the structural impact of the mutations on protein stability and provides strong support to the efficacy of the Potts model as an extensive “scanning” tool for mutational epistasis across the entire set of protein targets. Furthermore, the high correlation with FEP/MD on apo structures investigating mutational effects on stability indicates that the structural origins of the epistatic correlations involved in HIV drug resistance are features “intrinsic” to the protein conformational free energy landscape, there can be additional contributions as a consequence of drug binding. The findings suggest that HIV primarily leverages pre-existing intrinsic epistatic networks to explore resistance pathways that maximize fitness retention. Although drug selection pressure forces the virus to explore rugged regions of the fitness landscape that are otherwise poorly accessible, giving rise to DRMs, these mutations arise preferentially at strongly correlated sites and utilize the “intrinsic” network of epistatic interactions. Strongly coupled sites are more likely to be sites where clinically relevant DRMs arise because resistance mutations severely impair viral replication and are thus, unlikely to occur at independent or non-coupled sites where the virus doesn’t have viable pathways to remedy the fitness detriments caused by the mutation [4]. When the environment in which HIV replicates changes due to the initiation of drug therapy, HIV must mutate in ways that abrogate drug binding, while at the same time preserving protein function. Large epistatic interactions connect sites that can co-mutate with limited costs to fitness, even if the associated individual mutations are costly. Such sets of positions are therefore more likely to be associated with resistance. Here our assumption is that resistance cannot be achieved through selectively neutral mutations at single sites, in which case drug treatment would likely be ineffective.

### Structural underpinnings of the strongest epistatic interactions involving drug-resistance mutations

The structure-based analyses using FEP/MD illustrate the biophysical origins of the mutational correlations. In this section, we examine the structural rationale for the strongest epistatic couplings with a focus on HIV IN. Figure 3 shows Cα–Cα distances for the top 20 synergistically (blue-green) and antagonistically (red-orange) coupled residue pairs in HIV-1 PR, RT, and IN in WT apo structures. Residue pairs fall into two categories: (1) direct contacts, which include both adjacent residues (≤5 Å) and spatially close but non-adjacent residues (5–10 Å), and (2) indirect contacts, with distances >10 Å, likely reflecting interactions mediated long range electrostatic interactions [72] and/or allosteric effects. A clear trend across the three enzymes suggests that many of the strongest synergistic and antagonistic couplings occur between residues located within 5–10 Å, indicating that local structural interactions play a central role in shaping epistatic effects. In contrast to PR and RT, IN exhibits a broader distribution of residue pair distances, with a substantial number of strongly coupled residues located beyond 10 Å (in the 10–25 Å range) suggesting that epistatic effects in IN are more frequently mediated by long-range interactions, likely reflecting its multi-domain architecture involving the N-terminal (NTD), catalytic core (CCD), and C-terminal (CTD) domains as well as the involvement of the viral DNA. As an example, one of the top couplings involves inter-domain residue pairs such as 143-230 which is strongly synergistically coupled, indicating that mutations in one region can exert substantial effects on the inter-protomer structural rearrangement in IN. Notably, many of the strongest epistatic interactions in IN also involve mutations to charged amino-acid residue types that can exert their influences over longer distances [72]. Overall, the findings suggest a distinct structural coupling mechanism in IN (**Fig. 4A-B**), where local contacts, inter-protomer interfaces, as well as long-range interactions collectively contribute to the biophysical bases for the strongest mutational correlations.

**Fig 3:**
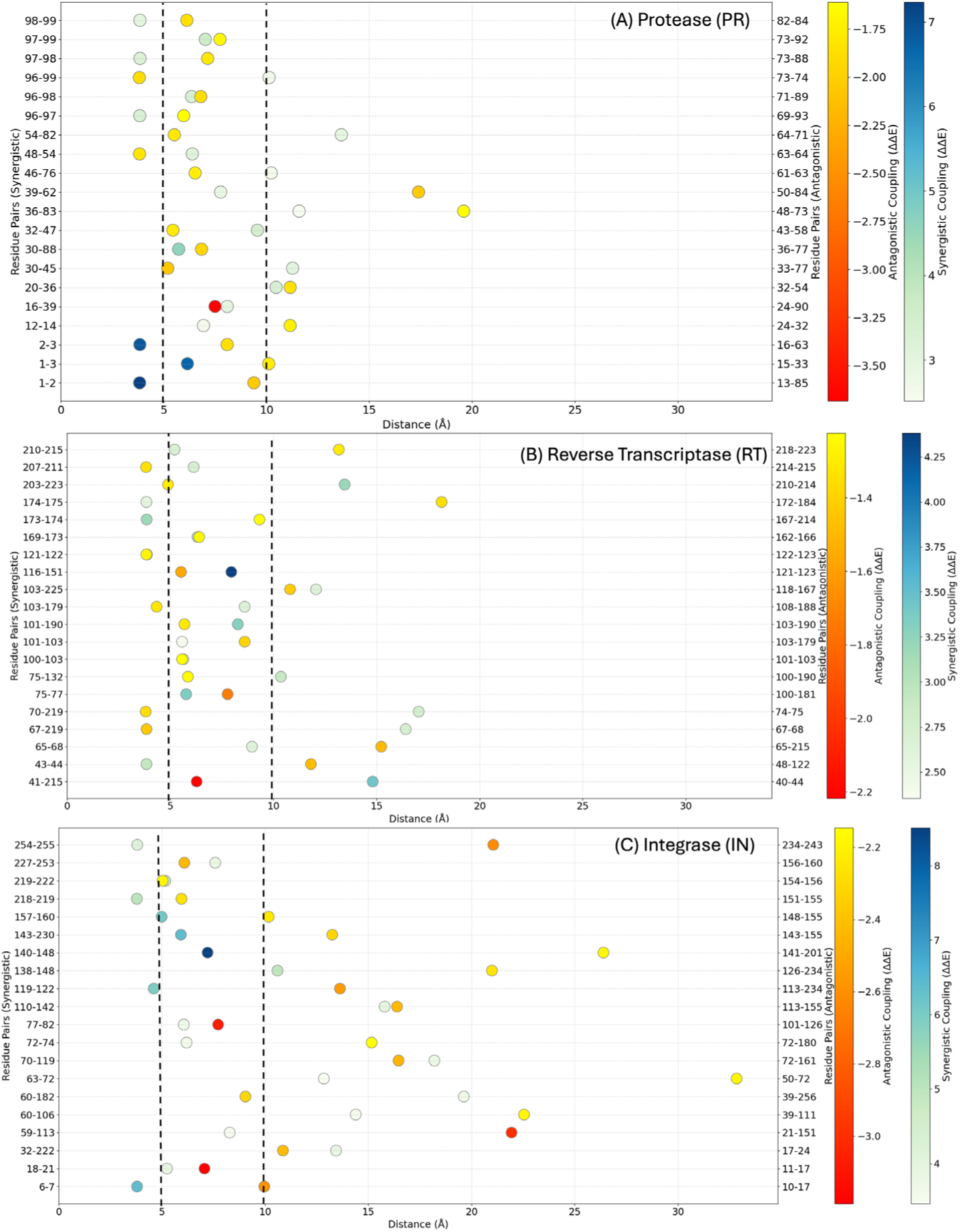
Fig 4: Structural overview of the strongest epistatic couplings between mutation pairs in HIV-1 (A) Protease (PR), (B) Reverse Transcriptase (RT) and (C) Integrase (IN) The plots show the distances between backbone carbon atoms (Cα) for the top 20 synergistically (shades of blue-green) and antagonistically (shades of red-yellow) coupled residue pairs in wild type apo structures of HIV-1 PR (PDB: 2PC0), RT (PDB: 1DLO) and IN (PDB:9C29). Both intra and inter-protomer distances between residue pairs are calculated and the shortest distance is reported. Only the strongest coupling between a pair of positions with multiple mutations in the top 20 are shown as the Cα-Cα distance remains same.

**Fig 4:**
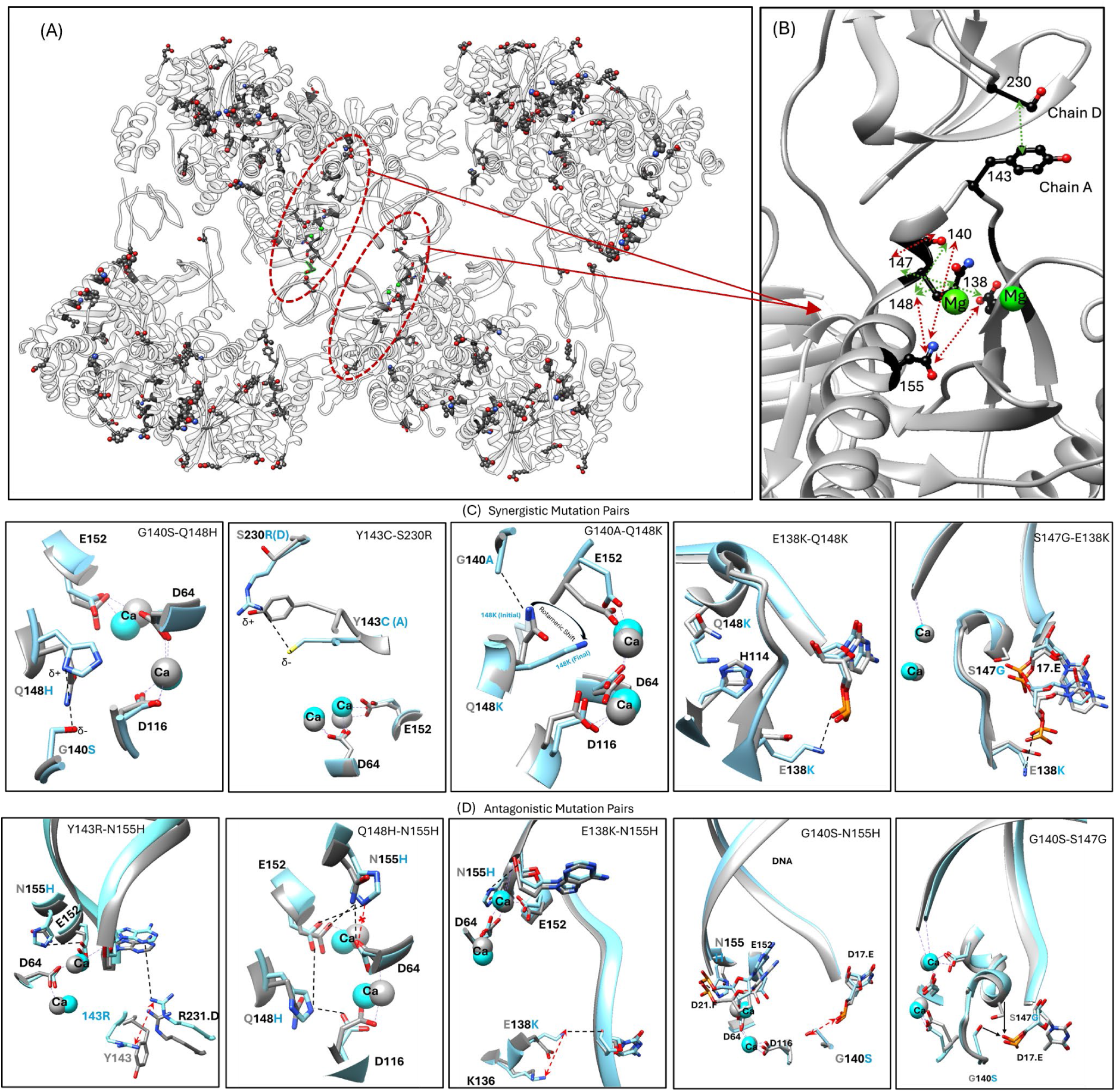
Structural Underpinnings of the strongest epistatic couplings between drug resistance associated mutation pairs in HIV-1 Integrase. (A) Residue positions of the top 40 strongly coupled mutation pairs (20 synergistic and 20 antagonistic) involving DRMs on the HIV-1 IN hexadecamer structure (9C29). The red dashed circle indicates the catalytic code domain (B) Close-up view of the IN catalytic core domain showing the top 10 strongly coupled DRM pairs (5 synergistic and 5 antagonistic). Green, black, blue and red color indicates Mg, C, N and O atoms respectively. **(C)** Panel shows the 5 synergistically coupled IN DRM pairs while (D) panel shows top 5 antagonistically coupled IN DRM pairs near the active site. For panels C and D the tetrameric HIV IN structure is used (6PUT). Light gray indicates WT and cyan color indicates the double mutant structures in C and D.

Next, we focus on the structural origins of the 10 strongest epistatic interactions in HIV IN that involve DRMs classified as ‘primary’ against the Integrase strand transfer inhibitors or INSTIs (**Fig 4)**. INSTIs form the first line of defense therapies in current ART [69, 70]. However, emerging DRMs have lowered the efficacy of even the latest INSTIs [70]. Probing how DRMs selected by INSTIs are epistatically linked to the structural level is essential to understanding the mechanisms in evolving drug resistance and revealing IN adaptations under selective pressure. The mutations are introduced into the apo IN structure and followed along MD trajectories (details on the generation of mutated Apo Integrase structures when not available from the PDB are provided in the Methods section).

To demonstrate this, we first focus on five highly synergistic IN DRM pairs selected based on the strength of DMC coupling and increasing prevalence in patients: G140A-Q148H, G140A-Q148K, Y143C-S230R, E138K-Q148K, and S147G-E138K (**Fig. 4C**). Full details and structural analyses are provided in the supplement (**Supplementary Information, Section S4)**.

Synergistic couplings arise when two disruptive mutations combine to restore or enhance function exerted by individual mutations through precise structural complementarity, often by forming new stabilizing interactions (**Supplementary Information, Table S2)** or enabling compensatory rearrangements that preserve catalytic loop and active site geometry. For example, in the **G140S-Q148H** DMC pair (ΔΔE = 8.5, δ = 4.5 ± 0.5 kcal/mol), Q148H alone introduces a charged imidazole into the catalytic core, destabilizing Mg²⁺ coordination that is involved in catalysis, while G140S increases loop polarity, rigidity and repulsive interaction with DNA. Both single mutations are disruptive to function[73-75]. However, when they arise together, Ser140 interacts with His148, neutralizing electrostatic strain and re-stabilizing the active site. Similar substitutions at position 140 like Thr or Cys would be too bulky or insufficiently electronegative, making the G140S-Q148H double mutation structurally optimal and the strongest synergistically coupled pair. Chemically similar mutations at position 148, such as 148R/K, also exhibit strong coupling with G140A/S (**Table 2A**). In the **G140A-Q148K** DMC pair (ΔΔE = 5.6, δ = 3.5 ± 0.78 kcal/mol), Q148K introduces a bulky, flexible lysine that disrupts loop conformation, misaligns the neighboring H114 residue, and like Q148H hinders enzyme catalysis [74]. G140A alone is also functionally disruptive[74]. The double mutant G140A-Q148K permits Lys148 to realign favorably with the Asp64-Asp116-Glu152 triad of residues involved in enzyme catalysis. In viral infectivity assays, the double mutant G140A-Q148K is epistatically compensatory in comparison to the additive effects of the two single mutations. The small size of alanine (and to a slightly lesser extent, Ser) is permissive, because bulkier substitutions would introduce significant steric constraints that are expected to compromise function. Q148R and Q148H similarly show strong synergistic epistatic coupling with G140A (**Table 2A**). In the **Y143C-S230R** DMC pair (ΔΔE = 6.5, δ = 3.93 ± 0.98 kcal/mol), individual mutations destabilize the inter-protomer interface. Cys143 loses the π-cation interaction with R231, and Arg230 introduces steric clash with Y143. However, together they are expected to re-stabilize the inter-protomer interface through mild ionic interactions (∼4 Å) that are optimal with this specific residue pairing. Other mutations that are known to arise at position 143[14], such as His, Lys, or Arg (Y143H/K/R) would repel Arg230, while Ala or Gly (Y143A/G) lack polarity for interaction. The higher electronegativity of Ser (Y143S) may cause an overly strong binding to Arg230, weakening synergy. Conversely, the size and charge to S230R explain why S230K also shows slightly weaker but still strong coupling with Y143C (ΔΔE = 3.03). In the **E138K-Q148K** DMC pair (ΔΔE = 5.0, δ = -3.29 ± 0.80 kcal/mol), E138K alone forms a new DNA contact that is mildly disruptive[74]. As indicated above, Q148K alone is a highly disruptive mutation because it destabilizes the catalytic loop by introducing a bulky positively charged group and also repositions His114. However, in combination, E138K offsets the rigidity introduced by Q148K and restores the orientation of His114. Like the G140A-Q148K double mutant, E138K-Q148K is epistatically compensatory in comparison to the additive effects of the two single mutations[74]. Finally, in the **E138K-S147G** DMC pair (ΔΔE = 2.6, vDNA strand[74], both of which are disruptive [76]. However, when present together, the two likely compensate for each other’s detrimental effects. Mutations such as E138A/T, which are observed in the Stanford HIVDB [14], will not form productive interactions with DNA that can be facilitated by S147G. The same would be expected of the Ala147 substitution, also observed in Stanford HIVDB[14], which may additionally locally rigidify the loop region surrounding the residue. Notably, the strongly synergistic focused mutation pairs resolve spatial or electrostatic conflicts that single mutations alone cannot accomplish, making these specific pairs structurally compatible and functionally compensatory in a way unmatched by alternative substitutions. In general, synergy does not solely depend on the proximity of residues but often arises from functional interdependence. These mutation pairs work in concert to create a more favorable structural or electrostatic environment that either could not achieve alone.

We also selected five highly antagonistic IN DRM mutation pairs (**Fig. 4D**) based on strong coupling via DMC analysis: Y143R-N155H, E138K-N155H, Q148H-N155H, G140S-N155H, and G140S-S147G. We note that N155H, which is a major and frequently observed DRM, is consistently paired with numerous residues to form strongly antagonistically coupled mutation pairs. Full details and structural analyses are provided in the supplement.

Antagonistic pairs combine mutations that individually impair structure or function and, when co-occurring, exacerbate these defects through conflicting spatial or electrostatic effects (**Supplementary Information, Table S2**). For **Y143R-N155H** DMC pair (ΔΔE = -2.31, δ = 3.06 ± 0.64 kcal/mol), individually, N155H introduces a new salt-bridge interaction with the vDNA strand, and it may also partially intrude on the Mg^2+^ chelation mediated by Asn155 and the catalytic residue Glu152. When paired with Y143R, the additional electrostatic repulsion mediated by Y143R interactions with Arg231 of an adjacent protomer misdirect DNA contacts and compound structural instability. Similarly, for **Q148H-N155H (Fig. 4a** DMC pair (ΔΔE = -2.24, δ = 3.47 ± 0.53 kcal/mol), both the Q148H and N155H single mutations heavily intrude into the active site, via introduction of bulky positive charges that disrupt the Asp64 and Glu152 chelation of the catalytic Mg^2+^ ions (in addition to affecting the configuration of the vDNA). Together, these detrimental interactions generate structural tension across the active site, creating a molecular tug-of-war, resulting in a non-additive, antagonistic outcome. Stronger antagonism with Q148R/K may stem from the shared chemical nature of H, R, and K residues (**Table 2B**). The **E138K-N155H** DMC pair (ΔΔE = -1.2, δ = 1.7 ± 0.89 kcal/mol), whose individual effects are described above, similarly suffer from rigidification of the vDNA via the introduction of two new salt-bridge interactions locally onto the same strand. Together, these mutations likely misalign active site flexibility via destabilization of DNA contacts that are necessary for catalysis. In the **G140S-N155H** DMC pair (ΔΔE = -2.12, δ = 2.19 ± 0.77 kcal/mol), the G140S and N155H mutations generate diverging electrostatic forces in the active site that physically strain enzyme activity, as well as the DNA–protein interface. Cys140, being less electronegative, fails to generate enough electrostatic force to put physical strain on the DNA backbone in comparison to Ser140, but the G140C-N155H pair remains antagonistically coupled (ΔΔE = -1.02). Lastly, in the **G140S-S147G** DMC pair (ΔΔE = -2.10, δ = 1.37 ± 0.47 kcal/mol), G140S introduces a partly negatively charged hydroxyl that increases loop polarity and rigidity and repulsive interaction with DNA, while S147G removes a stabilizing DNA contact, the combination of which led to significant local destabilization of the active site. Both Cys (ΔΔE = -1.95) and Thr (ΔΔE = -1.83) at position 140 show similar antagonistic behavior when coupled with S147G. Collectively, these antagonistic mutation pairs are especially detrimental, because their spatial footprints and chemical effects overlap or clash, with no possibility for mutual compensation. This makes them structurally incompatible and functionally antagonistic in comparison to other possible combinations (**Supplementary Information, Section S4**).

## Discussion

HIV mutates rapidly as it jumps from host to host, acquiring resistance to each host’s distinct immune response and applied drug regimen [36]; the rapid evolution of drug resistance in HIV remains a formidable challenge to the efficacy of ART [1-3]. When evolving under the selective pressure of ART, DRMs provide viable escape pathways for the virus and are accompanied by associated mutations, which compensate for the initial fitness penalty of incurring a primary resistance mutation [10-12]. Understanding epistatic interactions is essential for predicting which combinations of mutations enable HIV to maintain replication capacity while developing resistance to antiretroviral drugs under therapeutic selection pressure.

We present a framework that integrates sequence-based evolutionary modeling and structure-based molecular simulations to identify the strongest cooperative pairs of drug resistance mutations and explain the molecular basis for the epistatic correlations that govern resistance evolution. By leveraging a Potts model-based statistical approach, we systematically map the mutational fitness landscape of HIV’s three primary drug target enzymes—PR, RT, and IN. Our results reveal that the strongest epistatic interactions involve known drug resistance-associated mutations, underscoring the role of the intrinsic protein stability and fitness landscape as HIV evolves under drug selection pressure. Furthermore, the molecular dynamics-based free energy calculations provide direct structural and thermodynamic evidence that these correlated effects are largely driven by “intrinsic” effects on protein stability in contrast to correlations arising from interactions with the bound drug which are associated with non-linear affects on the binding affinity itself. Importantly, the very high degree of correlation between the thermodynamic signatures estimated by the Potts statistical energies with changes in viral protein stability for the HIV viral enzymes is unlike those observed in the literature for other eukaryotic and prokaryotic enzymes [24, 45]. *Zhang et al*. [25] observe lower correlation between the Potts model predicted epistasis and mutant protein stability effects predicted by FoldX or Rosetta for mutations at DRAPS in HIV-1 PR; both FoldX and Rosetta predict the stability effects of single mutations well, but are unable to account for epistatic effects that can significantly affect stability and become more pronounced for higher orders of mutations. The FEP/MD simulations presented in this work not only provide a quantitative measure of epistatic effects on protein stability, the high degree of correlation with thermodynamic signatures estimated by Potts statistical energies also suggests a special feature of viral evolution under drug selection pressure, indicating that drug selection pressure can introduce strong inductive biases modulating the evolutionary trajectories of viruses such as HIV into previously inaccessible regions of the fitness landscape while constraining resistance mutations preferentially emerge at pre-existing (in the absence of drug pressure) epistatically coupled sites [77], and reinforcing the idea that HIV leverages the intrinsic mutational landscapes to maintain fitness while circumventing therapeutic selection pressure.

Although drug selection pressure forces the virus to explore rugged regions of the fitness landscape otherwise not accessible, giving rise to resistance mutations, these mutations arise preferentially at strongly correlated sites and utilize the “intrinsic” network of epistasis. When the environment in which HIV replicates changes due to the initiation of drug therapy, HIV must mutate in ways that abrogate drug binding, while at the same time preserving protein function. Large epistatic interactions connect sites that are likely to be able to co-mutate with limited costs to fitness, even if the associated individual mutations are costly. Such sets of sites are therefore more likely to be associated with resistance. Here our assumption is that resistance cannot be achieved through selectively neutral mutations at single sites, in which case drug treatment would likely be ineffective. Drugs selecting for resistance mutations that occur without substantial fitness penalties in the wild-type virus, thus, have low barriers to resistance. Fitness detrimental resistance mutations, however, are unlikely to occur at non-interacting residue positions, due to a lack of viable opportunities to ameliorate the fitness defect [77]. Thus, HIV appears to exploit structurally constrained mutation networks, preferentially selecting resistance mutations that arise within pre-established cooperative frameworks.

The results suggest that major drug resistance sites can be identified based on the knowledge of the strongest epistatic interactions [4] from the DMC scans using the Potts statistical energy model, and applied to predict HIV evolution in response to new treatment regimens or vaccine candidates. We note, however, that this identification of resistance sites is “indirect” in the sense that only information about fitness is used. Thus, these predictions are not specific to a particular drug and would be most useful in cases when resistance information is unknown. In order to make highly accurate, drug-specific predictions of resistance, additional information would be required to narrow down the list of potential drug resistance sites identified based on fitness constraints to the ones that are most relevant for a particular case.

The insights also have profound implications for rational drug design and next-generation combination therapies. By identifying strongly coupled mutational networks, our framework enables the strategic design of inhibitors that exploit antagonistic epistatic relationships, potentially delaying or preventing the emergence of resistance. Targeting conserved, epistatically constrained regions may provide a more robust approach for future antiviral strategies. Moreover, by elucidating the structural and evolutionary basis of epistasis in HIV drug resistance, we provide a foundational step towards anticipating and mitigating resistance before it arises, paving the way for more durable and effective antiviral strategies. DRMs can be anticipated at residue positions that directly interact with the drug as well as have strong synergistic coupling with other residues to offer viable escape pathways through series of correlated mutations.

From a therapeutic perspective, these results highlight the potential of targeting antagonistically coupled sites to constrain the virus’s adaptive landscape. Designing inhibitors that destabilize resistance-prone epistatic networks or exploiting antagonistic interactions in c-ART could serve as viable strategies to extend drug efficacy and mitigate the emergence of resistance. Moving forward, these methodologies can be extended to predict HIV evolution under novel therapeutic pressures, providing a predictive framework for resistance pathways against emerging drugs and treatment regimens. Additionally, applying similar epistatic network analyses to other rapidly evolving pathogens, such as influenza or SARS-CoV-2, could offer transformative insights into the broader principles of drug resistance evolution.

Structural analysis (**Fig. 4C**) reveals synergistic couplings arise when two individually disruptive mutations complement each other’s structural or electrostatic deficiencies, resulting in restoration or enhancement of function. We exemplify this using the IN system, but the same principles should apply to other HIV enzymes. A recurring theme is that each mutation alone perturbs critical regions, such as loop conformation, active site charge balance, DNA interactions, etc., but when paired, they establish new stabilizing interactions (e.g., salt bridges, polar interaction, repositioned side chains) that resolve these defects. Notably, favorable outcomes to enzyme structure and function depend on precise chemical and geometric compatibility. For example, Ser140 uniquely facilitates a stabilizing salt bridge with His148, whereas Ala140 facilitates conformational flexibility that repositions Lys148 away from its location in the single Q148K substitution, which is otherwise harmful to the enzyme [74]. Some pairs, like E138K–Q148K or E138K–S147G, exhibit synergy despite not being in direct contact, and we presume that their functionally compensatory effect may involve local protein loop and DNA strand dynamics. Precise mechanisms would require determining high-resolution structures and detailed enzymatic analyses. These structural accommodations are unique towards specific residue combinations, making these coupled pairs not only cooperative but also evolutionarily favored due to their ability to preserve catalytic geometry and restore enzyme function.

In contrast, antagonistic couplings reflect structural incompatibility that often impose conflicting forces on the same domain or interface, amplifying destabilization rather than correcting it. Frequently, both mutations in an antagonistic pair affect the similar functional site or exert vector forces in opposite directions, leading to electrostatic clashes, loop strain, or DNA misalignment. For instance, N155H is a common antagonist due to its disruption of key salt bridges, in the viral DNA and in the active site. When paired with residues like Arg143, His148, or Ser147, the resulting interactions likely create competing DNA contacts or misorient loop geometries, which cannot be reconciled. These mutations fail to complement one another, leading to exacerbated structural defects that go beyond simply additive contributions, with significant loss of catalytic function.

Overall, synergistic mutation pairs demonstrate functional interdependence through precise structural compensation, while antagonistic pairs reveal mechanistic conflict, where overlapping or opposing effects disrupt protein stability. This framework underscores the importance of context-specific interaction networks where synergy depends not just on proximity, but on the ability of one mutation to restore what the other mutation disrupts, whereas antagonism results from redundancy, repulsion, or incompatible conformational changes.

## Methods

In this section, we present the computational models used in the study.

### Potts co-evolutionary model of protein fitness landscapes

We use the Potts model which is a probabilistic model that estimates the probability of observing specific protein sequences by capturing both individual amino acid frequencies and pairwise interactions between positions, constrained only to match the observed single-site and pairwise frequencies in the patient-derived sequence data [49, 78-80]. The Potts model has a long history of use in statistical physics and analysis of protein sequence. In a set of protein sequences, the single and pair amino acid frequencies are average quantities that can be estimated from the finite samples using the data.

To build the Potts inference, the goal is to approximate the unknown empirical probability distribution *P*(*S*) which best describes HIV-1 sequences *S* of length *L*, where each residue is encoded in an alphabet *Q*, by a model probability distribution *P*^*m*^(*S*) as in [81]. We choose the ‘least biased’ or maximum entropy distribution as the model distribution. Such distributions maximizing the entropy have been previously derived in [42, 43, 54, 79, 80] with the constraint that the empirical univariate and bivariate marginal distributions are effectively preserved. We follow a derivation of the maximum entropy model using Lagrange multipliers as in [43, 81]. Our maximum entropy model takes the form of an exponential distribution given by:

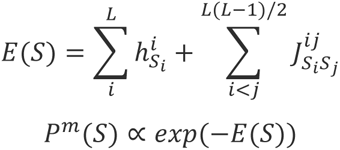

where the quantity *E*(*S*) is the Potts Hamiltonian determining the statistical energy of a sequence *S* of length *L*, the model parameters 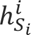 called ‘fields’ represent the statistical energy of residue *S*_*i*_ at position *i* in sequence *S*, and 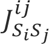 are ‘couplings’ representing the energy contribution of a position pair *i*, *j*. In this form, the Potts Hamiltonian consists of *LQ* field parameters 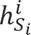 and 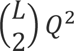 coupling parameters 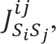 and for the exponential distribution *P*^*m*^(*S*) ∝ *exp*(−*E*(*S*), negative fields and couplings signify favored amino acids. The change in Potts energy due to mutating a residue *α* at position *i* in *S* to *β* is then given by:

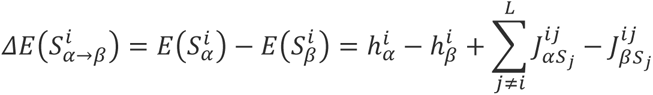

In this form, a 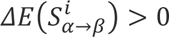 implies that the residue *β* is more favorable than residue *α* at position *i* for the given sequence *S*. If *α* represents the wild-type residue at *i* and *β* the mutant, then the mutant is favorable over the wild-type if *ΔE* > 0 for the change, and *vice versa*. Further details of the models are described in our previous work [6, 18, 82]. The sample size of the MSA plays a critical role in determining the quality and effectiveness of the model [83] and we confirm that the models are fit using sufficient data with minimal overfitting.

### Data processing for the Potts model, mutation classification, and mutation overlap under different drugs

We collect sequence, patient as well as reference information from the Stanford HIV HIVDB (https://hivdb.stanford.edu) [15, 84]. In this work, we have used the Stanford HIVDB genotype-rx (https://hivdb.stanford.edu/pages/genotype-rx.html) to obtain protein sequences for HIV target proteins. Alternatively, downloadable sequence datasets are also available on the Stanford HIVDB (https://hivdb.stanford.edu/pages/geno-rx-datasets.html). The filtering criteria we used are: HIV-1, subtype B and nonCRF, and drug-experienced (# of PI = 1–9 for PR, # of NRTI = 1–9 and # of NNRTI = 1–4 for RT, and # of INST = 1–3 for IN), removal of mixtures, and unambiguous amino acid sequences (amino acids are ‘-ACDEFGHIKLMNPQRSTVWY’). For RT, sequences with exposure to both NRTIs and NNRTIs were selected. Sequences with insertions (‘#’) and deletions (‘∼’) are removed. MSA columns and rows with more than 1% gaps (‘.’) are removed. To retain enough sequence coverage in the MSA, we removed residues: residues 1–38 and residue 227 onwards for RT, and 264 onwards for IN. For this reason, some interesting DRMs like F227I/L/V/C, L234I, P236L or N348I (NNRTI affected) for RT are not amenable for our analysis. The protein sequences for the molecular clones NL4-3 and HXB2 are obtained from GenBank [85] with accession number AF324493.2 and K03455.1, respectively. The protein ids for the pol polyprotein are AAK08484.2 and AAB50259.1, respectively. The subtype B consensus sequence is obtained from the Los Alamos HIV sequence database [86] consensus and ancestral sequence alignments (https://www.hiv.lanl.gov/content/sequence/HIV/CONSENSUS/Consensus.html, last updated August 2004), and is also available in Appendix 1 of the Stanford HIVDB release notes page (https://hivdb.stanford.edu/page/release-notes/). Sequences are given weights reciprocal to the number of sequences contributed by each patient such that if multiple sequences are obtained from a single patient, the effective number of sequences obtained from any single patient is still 1. Sequences from different patients are assumed to be independent. It has been shown that phylogenetic trees of drug-naive and drug-experienced HIV-1 patients exhibit star-like phylogenies [87, 88], and thus, phylogenetic corrections are not required. Potts models of different HIV-1 protein sequences under immune as well as drug selection pressure have also previously been parameterized without phylogenetic corrections [4, 6, 7, 18, 20, 21, 89, 90].

In this work, mutations in the HIV genome have been classified into three main classes: primary or signature drug-resistance mutations, accessory or secondary mutations, and polymorphic mutations. The subtype B consensus sequence, which is derived from an alignment of subtype B sequences maintained at the Los Alamos HIV Sequence Database (https://www.hiv.lanl.gov), and is a commonly used reference sequence to which new sequences are compared, is used as our reference sequence and is referred to as the ‘consensus wild-type’ throughout the text. In accordance with current literature [66, 91] and the Stanford HIVDB, any mutation that (i) affects in vitro drug-susceptibility, (ii) occurs commonly in persons experiencing virological failure, and (iii) occurs with fairly low extent of polymorphism (mutations occurring commonly as natural variants) among untreated individuals, is classified as a major or primary drug-resistance mutation. In contrast, mutations with little or no effect on drug susceptibility are classified as accessory or secondary. Such mutations may reduce drug susceptibility or increase replication fitness only in combination with a primary mutation. Polymorphic mutations are mutations occurring as natural variants typically in antiretroviral drug (ARV) naive patients. Polymorphic mutations may occur in the absence of drug pressure and usually have little effect on ARV susceptibility when they occur without other DRMs. In this regard, polymorphic mutations affecting ARV susceptibility in combination with other DRMs can be classified as accessory.

To classify the overlap of mutations selected by different ART drugs within the same class, we used the frequency of drug resistance mutations at each residue position in sequences labelled with particular drug usage in the Stanford HIVDB similar to [6] (Fig 8 in [6]), Stanford HIVDB Drug Resistance Summaries, and the latest prevalence-based plots in [91]. Most mutations that occur in response to one drug are largely observed to have occurred when treated with another drug of the same class, restricting the choice of DRM pairs where each individual DRM is largely selected by one (set of) drug(s) and not by another (set of) drug(s) within the same class. The exception to this general rule are specific patterns of mutations selected by the NRTIs (Thymidine Analog mutations or TAMs, and the generally TAM-exclusive Q151M complex) [66, 91]. When choosing for DRM pairs where each individual DRM is selected by a different class of drugs, we have limited our scope to only the NRTI and NNRTI classes of RT inhibitors, and to the top 20 strongest observed interactions through Potts model double mutant cycles. Interactions between mutations in drug-target proteins such as PR and IN or PR and RT are not amenable to our analyses through models built separately based on individual drug target protein sequences.

### Alchemical Free Energy Perturbation (FEP) Simulation

Simulations of the alchemical free energy protocol were carried out in GROMACS 2023 [92, 93] for HIV PR, RT, and IN. The HIV mutants were created starting from the WT apo structures of PR (PDB:2PC0) [94], RT (PDB:1DLO) [95] and IN a (PDB:6PUT) [96]. using the pmx tool [97] which is able to generate hybrid structures and topologies for amino acid mutations. Each protein was placed in a TIP3P [98] triclinic water box. For HIV IN, NaCl ions [99] were added to neutralize the system and reach concentration of 0.15 mol/L. The force field of choice was AMBER99sb*ILDN [100, 101] which provided accurate results in previous alchemical free energy studies [102].

All the simulations were performed at 300 K temperature using the stochastic velocity rescaling thermostat [103] with a time constant of 1.0 ps. The pressure was kept at 1 bar using the Parrinello-Rahman barostat [104] with the time constant of 5 ps. The electrostatic interactions were treated by means of PME [105] with the real space cutoff of 1.0 nm and the grid spacing of 0.12 nm. The van der Waals interactions were shifted to zero at the cutoff of 1.0 nm with energy and pressure dispersion corrections applied analytically. An analytical dispersion correction for the energy and pressure was applied. For alchemical free energy calculations, soft-core potentials were applied to handle non-bonded interactions smoothly across λ states. A total of 27 λ windows were used, spanning from 0 to 1 with finer resolution near the end states. The production simulations were carried out for 25 ns for each λ window, with a time step of 2 fs. Free energy differences (ΔG) between neighboring λ states were calculated using the Bennett’s Acceptance Ratio (BAR) [106] method as implemented in GROMACS. This approach estimates the free energy change by statistically reweighting energy differences between adjacent alchemical states, providing a robust and accurate estimate of the overall free energy profile across the λ windows. To improve sampling across alchemical states, λ-Replica Exchange Molecular Dynamics (λ-REMD) [107] was employed for a few mutations pairs specifically in IN. In this approach, 26 replicas were simulated in parallel, each corresponding to a different value along the alchemical λ coordinate. Periodic exchange attempts were carried out between neighboring λ states based on the Metropolis criterion, allowing configurations to traverse λ space and escape local minima. This method enhances the efficiency and convergence of free energy calculations by promoting better overlap between adjacent alchemical states and reducing hysteresis in forward and reverse transitions. The calculated and experimentally measured values were compared by spearman correlation.

### Molecular Dynamics Simulations

An additional classical molecular dynamics simulation was carried out to further investigate the structural underpinnings of the coupled mutations. The final frame (FEP simulation) from the λ = 1.0 state of the alchemical simulation was extracted, and the hybrid residue was converted to the fully mutated side chain. This modified structure was then used as the starting point for a 100 ns unbiased MD simulation in GROMACS [93] employing the AMBER99SB*-ILDN force field [100, 101]. The system was solvated in a TIP3P water box and neutralized with counterions. Temperature and pressure were maintained at 300 K and 1 bar, respectively, using the velocity-rescaling thermostat and the Parrinello–Rahman barostat [103]..

### Error Estimation via Bootstrapping and BAR Analysis

To estimate uncertainties in the calculated free energies, we employed a bootstrapping-based resampling approach followed by Bennett’s Acceptance Ratio (BAR) [106] analysis. The workflow is divided into two key stages: (1) generating resampled datasets and (2) extracting error estimates.

First, the trajectory data from each transformation window was divided into equal-sized blocks. These blocks were then randomly resampled with replacement to generate a specified number of bootstrap datasets. Each resampled dataset was saved as a new input file, providing multiple statistically independent inputs for subsequent analysis. This method captures the intrinsic variability within the simulation data and allows for robust estimation of statistical uncertainties.

Next, GROMACS was used to process the bootstrap-generated datasets and compute the free energy differences (ΔG) using BAR analysis. For each transformation, ΔG values across bootstrap samples were extracted and averaged. The overall uncertainty was computed via the root-sum-of-squares (RSS) method, capturing the propagated errors across all transformation points. Finally, the free energy results were converted from kJ/mol to kcal/mol for standard reporting.

This two-step bootstrapping and BAR analysis framework provides a statistically rigorous estimate of the mean and error bars, improving the confidence in reported ΔG values and allowing meaningful comparisons across mutational states.

### Ensemble modelling of structures for structural underpinnings of epistatic DRMs

The structural interaction analysis for DRMs used structures from experimental cryo-EM datasets of the oligomeric states of the proteins (hexadecamer PDB-9C29, EMD-45151), with additional conformers generated using a modified AlphaFold3 (AF3) [108-110] workflow. The modified AF3 deliberately reduces the sequence diversity by sub-sampling the sequence in the multiple sequence alignment (MSA) generated by AF3, enhancing conformational heterogeneity by reducing sequence/structure bias inherent in the AF3 training dataset. These sub-MSAs (sMSA) were generated by randomly selecting a small number (*n*=5, 10, 15) of sequences, which are then used for structure prediction. The resulting structural models were filtered and validated by a rigid body fit into the cryo-EM density maps to ensure conformational consistency with the experimental models.

## Supplementary Information

The supplementary information is available.

## Acknowledgement

This work has been supported by the National Institutes of Health through grants U54-AI150472, U54-AI170855, U01 AI136680, and R01 AI178849 (to D.L. and R.M.L.), and U54AI170792 (to K.H, A.S, and I.E), the Margaret T. Morris and the Hearst Foundations (to D.L.), R35-GM132090, and S10OD020095 (to R.M.L.), and California HIV Research Program H25TC9308 to I.C. The National Science Foundation also provided funding through a grant awarded to R.M.L and A.H (1934848). A.B was supported by the Eric and Wendy Schmidt AI in Science Postdoctoral Fellowship, a program of Schmidt Sciences. Computing resources were generously supported through an ACCESS proposal MCB100145 (R.M.L) and BIO240258 (D.L). Some of the computing core infrastructure was also supported by NIH-NCI CCSG P30 CA014195. The founders had no role in study design, data collection and analysis, decision to publish, or preparation of the manuscript.

## References

1. Ghosh, A.K., Four decades of continuing innovations in the development of antiretroviral therapy for HIV/AIDS: Progress to date and future challenges. Global Health & Medicine, 2023. 5(4): p. 194–198.

2. Temereanca, A. and S. Ruta, Strategies to overcome HIV drug resistance-current and future perspectives. Frontiers in Microbiology, 2023. 14: p. 1133407.

3. Apetroaei, M.-M., et al., The Phenomenon of Antiretroviral Drug Resistance in the Context of Human Immunodeficiency Virus Treatment: Dynamic and Ever Evolving Subject Matter. Biomedicines, 2024. 12(4): p. 915.

4. Butler, T.C., et al., Identification of drug resistance mutations in HIV from constraints on natural evolution. Physical Review E, 2016. 93(2): p. 022412.

5. Flynn, W.F., et al., Inference of Epistatic Effects Leading to Entrenchment and Drug Resistance in HIV-1 Protease. Molecular Biology and Evolution, 2017. 34(6): p. 1291–1306.

6. Biswas, A., et al., Epistasis and entrenchment of drug resistance in HIV-1 subtype B. Elife, 2019. 8.

7. Biswas, A., A. Haldane, and R.M. Levy, Limits to detecting epistasis in the fitness landscape of HIV. PLOS ONE, 2022. 17(1): p. e0262314.

8. Poelwijk, F.J., M. Socolich, and R. Ranganathan, Learning the pattern of epistasis linking genotype and phenotype in a protein. Nature communications, 2019. 10(1): p. 4213.

9. Starr, T.N. and J.W. Thornton, Epistasis in protein evolution. Protein science, 2016. 25(7): p. 1204–1218.

10. Duan, S., et al., Epistatic interactions between neuraminidase mutations facilitated the emergence of the oseltamivir-resistant H1N1 influenza viruses. Nature Communications, 2014. 5(1): p. 5029.

11. Zhang, H., A.A. Quadeer, and M.R. McKay, Direct-acting antiviral resistance of Hepatitis C virus is promoted by epistasis. Nature communications, 2023. 14(1): p. 7457.

12. Heeney, J.L., A.G. Dalgleish, and R.A. Weiss, Origins of HIV and the evolution of resistance to AIDS. Science, 2006. 313(5786): p. 462-466.

13. Rhee, S.Y., et al., Human immunodeficiency virus reverse transcriptase and protease sequence database. Nucleic Acids Res, 2003. 31(1): p. 298–303.

14. Stnaford. Stanford HIV Database. September 15, 2022]; Available from: https://hivdb.stanford.edu/.

15. Shafer, R.W., Rationale and uses of a public HIV drug-resistance database. The Journal of infectious diseases, 2006. 194(Supplement_1): p. S51-S58.

16. Miton, C.M., K. Buda, and N. Tokuriki, Epistasis and intramolecular networks in protein evolution. Current opinion in structural biology, 2021. 69: p. 160–168.

17. Carter, P.J., et al., The use of double mutants to detect structural changes in the active site of the tyrosyl-tRNA synthetase (Bacillus stearothermophilus). Cell, 1984. 38(3): p. 835–840.

18. Choudhuri, I., et al., Contingency and Entrenchment of Drug-Resistance Mutations in HIV Viral Proteins. J. Phys. Chem. B, 2022.

19. Biswas, A., et al., Kinetic coevolutionary models predict the temporal emergence of HIV-1 resistance mutations under drug selection pressure. Proceedings of the National Academy of Sciences, 2024. 121(15): p. e2316662121.

20. Shekhar, K., et al., Spin models inferred from patient-derived viral sequence data faithfully describe HIV fitness landscapes. Physical review E, 2013. 88(6): p. 062705.

21. Mann, J.K., et al., The fitness landscape of HIV-1 gag: advanced modeling approaches and validation of model predictions by in vitro testing. PLoS computational biology, 2014. 10(8): p. e1003776.

22. Louie, R.H., et al., Fitness landscape of the human immunodeficiency virus envelope protein that is targeted by antibodies. Proceedings of the National Academy of Sciences, 2018. 115(4): p. E564–E573.

23. Quadeer, A.A., et al., Deconvolving mutational patterns of poliovirus outbreaks reveals its intrinsic fitness landscape. Nature Communications, 2020. 11(1): p. 377.

24. Riesselman, A.J., J.B. Ingraham, and D.S. Marks, Deep generative models of genetic variation capture the effects of mutations. Nature Methods, 2018. 15(10): p. 816–822.

25. Zhang, T.-h., et al., Predominance of positive epistasis among drug resistance-associated mutations in HIV-1 protease. PLoS genetics, 2020. 16(10): p. e1009009.

26. Gregson, J., et al., Global epidemiology of drug resistance after failure of WHO recommended first-line regimens for adult HIV-1 infection: a multicentre retrospective cohort study. The Lancet infectious diseases, 2016. 16(5): p. 565–575.

27. Posfai, A., et al., Selection for protein stability enriches for epistatic interactions. Genes, 2018. 9(9): p. 423.

28. Liberles, D.A. and A.I. Teufel, Evolution and structure of proteins and proteomes. 2018, MDPI. p. 583.

29. Duan, J., D. Lupyan, and L. Wang, Improving the accuracy of protein thermostability predictions for single point mutations. Biophysical Journal, 2020. 119(1): p. 115–127.

30. Ono, F., et al., Improvement in predicting drug sensitivity changes associated with protein mutations using a molecular dynamics based alchemical mutation method. Scientific Reports, 2020. 10(1): p. 2161.

31. Sergeeva, A.P., et al., Free energy perturbation calculations of mutation effects on SARS-CoV-2 RBD:: ACE2 binding affinity. Journal of Molecular Biology, 2023. 435(15): p. 168187.

32. Werner, M., V. Gapsys, and B.L. de Groot, One Plus One Makes Three: Triangular Coupling of Correlated Amino Acid Mutations. Journal of Physical Chemistry Letters, 2021. 12(12): p. 3195–3201.

33. Gizzio, J., et al., Evolutionary divergence in the conformational landscapes of tyrosine vs serine/threonine kinases. Elife, 2022. 11: p. e83368.

34. Gizzio, J., et al., Evolutionary sequence and structural basis for the distinct conformational landscapes of Tyr and Ser/Thr kinases. Nature Communications, 2024. 15(1): p. 6545.

35. Cuevas, J.M., et al., Extremely high mutation rate of HIV-1 in vivo. PLoS biology, 2015. 13(9): p. e1002251.

36. Coffin, J.M., HIV Population Dynamics in Vivo: Implications for Genetic Variation, Pathogenesis, and Therapy. Science, 1995. 267(5197): p. 483-489.

37. Perelson, A.S., et al., HIV-1 dynamics in vivo: virion clearance rate, infected cell life-span, and viral generation time. Science, 1996. 271(5255): p. 1582-1586.

38. Sever, B., et al., A review of FDA-approved anti-HIV-1 drugs, anti-gag compounds, and potential strategies for HIV-1 eradication. International Journal of Molecular Sciences, 2024. 25(7): p. 3659.

39. Troyer, R.M., et al., Variable fitness impact of HIV-1 escape mutations to cytotoxic T lymphocyte (CTL) response. PLoS pathogens, 2009. 5(4): p. e1000365.

40. Da Silva, J., et al., Fitness epistasis and constraints on adaptation in a human immunodeficiency virus type 1 protein region. Genetics, 2010. 185(1): p. 293–303.

41. Liu, Y., et al., A sensitive real-time PCR based assay to estimate the impact of amino acid substitutions on the competitive replication fitness of human immunodeficiency virus type 1 in cell culture. Journal of virological methods, 2013. 189(1): p. 157–166.

42. Morcos, F., et al., Direct-coupling analysis of residue coevolution captures native contacts across many protein families. Proc Natl Acad Sci U S A, 2011. 108(49): p. E1293–301.

43. Ferguson, A.L., et al., Translating HIV sequences into quantitative fitness landscapes predicts viral vulnerabilities for rational immunogen design. Immunity, 2013. 38(3): p. 606–617.

44. Jacquin, H., et al., Benchmarking inverse statistical approaches for protein structure and design with exactly solvable models. PLoS computational biology, 2016. 12(5): p. e1004889.

45. Hopf, T.A., et al., Mutation effects predicted from sequence co-variation. Nature biotechnology, 2017. 35(2): p. 128–135.

46. Morcos, F., et al., Direct coupling analysis for protein contact prediction. Methods Mol Biol, 2014. 1137: p. 55–70.

47. Morcos, F., et al., Coevolutionary information, protein folding landscapes, and the thermodynamics of natural selection. Proc Natl Acad Sci U S A, 2014. 111(34): p. 12408–13.

48. Marks, D.S., T.A. Hopf, and C. Sander, Protein structure prediction from sequence variation. Nature biotechnology, 2012. 30(11): p. 1072–1080.

49. Sułkowska, J.I., et al., Genomics-aided structure prediction. Proceedings of the National Academy of Sciences, 2012. 109(26): p. 10340–10345.

50. Sutto, L., et al., From residue coevolution to protein conformational ensembles and functional dynamics. Proceedings of the National Academy of Sciences, 2015. 112(44): p. 13567–13572.

51. Sjodt, M., et al., Structure of the peptidoglycan polymerase RodA resolved by evolutionary coupling analysis. Nature, 2018. 556(7699): p. 118-121.

52. Haldane, A., et al., Structural propensities of kinase family proteins from a Potts model of residue co-variation. Protein Sci, 2016. 25(8): p. 1378–84.

53. Haldane, A., et al., Coevolutionary Landscape of Kinase Family Proteins: Sequence Probabilities and Functional Motifs. Biophys J, 2018. 114(1): p. 21–31.

54. Barton, J.P., et al., ACE: adaptive cluster expansion for maximum entropy graphical model inference. Bioinformatics, 2016. 32(20): p. 3089–3097.

55. Levy, R.M., A. Haldane, and W.F. Flynn, Potts Hamiltonian models of protein co-variation, free energy landscapes, and evolutionary fitness. Current Opinion in Structural Biology, 2017. 43: p. 55–62.

56. Li, M., et al., Mechanisms of HIV-1 Integrase Resistance to Dolutegravir and Potent Inhibition of Drug Resistant Variants. bioRxiv, 2022.

57. Rhee, S.-Y., et al., HIV-1 protease mutations and protease inhibitor cross-resistance. Antimicrobial agents and chemotherapy, 2010. 54(10): p. 4253–4261.

58. Götte, M., et al., The M184V mutation in the reverse transcriptase of human immunodeficiency virus type 1 impairs rescue of chain-terminated DNA synthesis. Journal of virology, 2000. 74(8): p. 3579–3585.

59. Sarafianos, S.G., et al., Structures of HIV-1 reverse transcriptase with pre-and post-translocation AZTMP-terminated DNA. The EMBO journal, 2002.

60. Larder, B.A., S.D. Kemp, and P.R. Harrigan, Potential mechanism for sustained antiretroviral efficacy of AZT-3TC combination therapy. Science, 1995. 269(5224): p. 696-699.

61. Marcelin, A.-G., Resistance to nucleoside reverse transcriptase inhibitors. Antiretroviral resistance in clinical practice, 2006.

62. Das, K., et al., Structural basis for the role of the K65R mutation in HIV-1 reverse transcriptase polymerization, excision antagonism, and tenofovir resistance. Journal of Biological Chemistry, 2009. 284(50): p. 35092–35100.

63. Parikh, U.M., et al., The K65R mutation in human immunodeficiency virus type 1 reverse transcriptase exhibits bidirectional phenotypic antagonism with thymidine analog mutations. Journal of virology, 2006. 80(10): p. 4971–4977.

64. Garforth, S.J., C. Lwatula, and V.R. Prasad, The lysine 65 residue in HIV-1 reverse transcriptase function and in nucleoside analog drug resistance. Viruses, 2014. 6(10): p. 4080–4094.

65. White, K.L., et al., The K65R reverse transcriptase mutation in HIV-1 reverses the excision phenotype of zidovudine resistance mutations. Antiviral therapy, 2006. 11(2): p. 155–163.

66. Wensing, A.M., et al., 2019 update of the drug resistance mutations in HIV-1. Top Antivir Med, 2019. 27(3): p. 111-121.

67. Izopet, J., et al., Shift in HIV resistance genotype after treatment interruption and short-term antiviral effect following a new salvage regimen. Aids, 2000. 14(15): p. 2247–2255.

68. Yang, W.-L., et al., Persistence of transmitted HIV-1 drug resistance mutations associated with fitness costs and viral genetic backgrounds. PLoS pathogens, 2015. 11(3): p. e1004722.

69. Gandhi, R.T., et al., Progressive reversion of human immunodeficiency virus type 1 resistance mutations in vivo after transmission of a multiply drug-resistant virus. Clinical infectious diseases, 2003. 37(12): p. 1693–1698.

70. Werner, M., V. Gapsys, and B.L. de Groot, One Plus One Makes Three: Triangular Coupling of Correlated Amino Acid Mutations. J Phys Chem Lett, 2021. 12(12): p. 3195–3201.

71. Gizzio, J., et al., Evolutionary sequence and structural basis for the distinct conformational landscapes of Tyr and Ser/Thr kinases. Nat Commun, 2024. 15(1): p. 6545.

72. Haq, O., et al., Correlated electrostatic mutations provide a reservoir of stability in HIV protease. 2012.

73. Malet, I., et al., Mutations associated with failure of raltegravir treatment affect integrase sensitivity to the inhibitor in vitro. Antimicrob Agents Chemother, 2008. 52(4): p. 1351–8.

74. Li, M., et al., Mechanisms of HIV-1 integrase resistance to dolutegravir and potent inhibition of drug-resistant variants. Sci Adv, 2023. 9(29): p. eadg5953.

75. Cook, N.J., et al., Structural basis of second-generation HIV integrase inhibitor action and viral resistance. Science, 2020. 367(6479): p. 806-810.

76. Andreatta, K., et al., Integrase inhibitor resistance selections initiated with drug resistant HIV-1. CROI, Boston, USA, 2018.

77. Shi, J., et al., Compensatory substitutions in the HIV-1 capsid reduce the fitness cost associated with resistance to a capsid-targeting small-molecule inhibitor. J Virol, 2015. 89(1): p. 208–19.

78. Cocco, S., et al., Inverse statistical physics of protein sequences: a key issues review. Reports on Progress in Physics, 2018. 81(3): p. 032601.

79. Mézard, M. and T. Mora, Constraint satisfaction problems and neural networks: A statistical physics perspective. Journal of Physiology-Paris, 2009. 103(1-2): p. 107–113.

80. Weigt, M., et al., Identification of direct residue contacts in protein–protein interaction by message passing. Proceedings of the National Academy of Sciences, 2009. 106(1): p. 67–72.

81. Mora, T. and W. Bialek, Are biological systems poised at criticality? Journal of Statistical Physics, 2011. 144: p. 268–302.

82. Haldane, A. and R.M. Levy, Mi3-GPU: MCMC-based inverse Ising inference on GPUs for protein covariation analysis. Computer Physics Communications, 2021. 260: p. 107312.

83. Haldane, A. and R.M. Levy, Influence of multiple-sequence-alignment depth on Potts statistical models of protein covariation. Phys Rev E, 2019. 99(3-1): p. 032405.

84. Rhee, S.-Y., et al., Human immunodeficiency virus reverse transcriptase and protease sequence database. Nucleic acids research, 2003. 31(1): p. 298–303.

85. Clark, K., et al., GenBank. Nucleic acids research, 2016. 44(D1): p. D67–D72.

86. Compendium, H.S., Foley B, Leitner T, Apetrei C, Hahn B, Mizrachi I, Mullins J, Rambaut A, Wolinsky S, and Korber B, Eds. Published by Theoretical Biology and Biophysics Group, Los Alamos National Laboratory, NM, LA-UR, 2013: p. 13-26007.

87. Keele, B.F., et al., Identification and characterization of transmitted and early founder virus envelopes in primary HIV-1 infection. Proceedings of the National Academy of Sciences, 2008. 105(21): p. 7552–7557.

88. Gupta, A. and C. Adami, Strong selection significantly increases epistatic interactions in the long-term evolution of a protein. PLoS genetics, 2016. 12(3): p. e1005960.

89. Flynn, W.F., et al., Deep Sequencing of Protease Inhibitor Resistant HIV Patient Isolates Reveals Patterns of Correlated Mutations in Gag and Protease. PLOS Computational Biology, 2015. 11(4): p. e1004249.

90. Li, M., et al., Mechanisms of HIV-1 integrase resistance to dolutegravir and potent inhibition of drug-resistant variants. Science Advances. 9(29): p. eadg5953.

91. Wensing, A.M., et al., 2022 update of the drug resistance mutations in HIV-1. Topics in antiviral medicine, 2022. 30(4): p. 559.

92. Berendsen, H.J., D. van der Spoel, and R. van Drunen, GROMACS: A message-passing parallel molecular dynamics implementation. Computer physics communications, 1995. 91(1-3): p. 43–56.

93. Van Der Spoel, D., et al., GROMACS: fast, flexible, and free. Journal of computational chemistry, 2005. 26(16): p. 1701–1718.

94. Heaslet, H., et al., Conformational flexibility in the flap domains of ligand-free HIV protease. Biological Crystallography, 2007. 63(8): p. 866–875.

95. Hsiou, Y., et al., Structure of unliganded HIV-1 reverse transcriptase at 2.7 Å resolution: implications of conformational changes for polymerization and inhibition mechanisms. Structure, 1996. 4(7): p. 853–860.

96. Passos, D.O., et al., Structural basis for strand-transfer inhibitor binding to HIV intasomes. Science, 2020. 367(6479): p. 810-814.

97. Gapsys, V., et al., Calculation of binding free energies. Molecular Modeling of Proteins, 2015: p. 173–209.

98. Jorgensen, W.L., et al., Comparison of simple potential functions for simulating liquid water. The Journal of chemical physics, 1983. 79(2): p. 926–935.

99. Joung, I.S. and T.E. Cheatham III, Determination of alkali and halide monovalent ion parameters for use in explicitly solvated biomolecular simulations. The journal of physical chemistry B, 2008. 112(30): p. 9020–9041.

100. Hornak, V., et al., Comparison of multiple Amber force fields and development of improved protein backbone parameters. Proteins: Structure, Function, and Bioinformatics, 2006. 65(3): p. 712–725.

101. Lindorff-Larsen, K., et al., Improved side-chain torsion potentials for the Amber ff99SB protein force field. Proteins: Structure, Function, and Bioinformatics, 2010. 78(8): p. 1950–1958.

102. Gapsys, V., et al., Accurate and rigorous prediction of the changes in protein free energies in a large-scale mutation scan. Angewandte Chemie International Edition, 2016. 55(26): p. 7364–7368.

103. Bussi, G., D. Donadio, and M. Parrinello, Canonical sampling through velocity rescaling. The Journal of chemical physics, 2007. 126(1).

104. Parrinello, M. and A. Rahman, Polymorphic transitions in single crystals: A new molecular dynamics method. Journal of Applied physics, 1981. 52(12): p. 7182–7190.

105. Darden, T., D. York, and L. Pedersen, Particle mesh Ewald: An N log (N) method for Ewald sums in large systems. Journal of chemical physics, 1993. 98: p. 10089–10089.

106. Bennett, C.H., Efficient estimation of free energy differences from Monte Carlo data. Journal of Computational Physics, 1976. 22(2): p. 245–268.

107. Qi, R., et al., Replica exchange molecular dynamics: a practical application protocol with solutions to common problems and a peptide aggregation and self-assembly example. Peptide self-assembly: Methods and protocols, 2018: p. 101–119.

108. Abramson, J., et al., Accurate structure prediction of biomolecular interactions with AlphaFold 3. Nature, 2024. 630(8016): p. 493-500.

109. Desai, D., et al., Review of AlphaFold 3: transformative advances in drug design and therapeutics. Cureus, 2024. 16(7).

110. Wayment-Steele, H.K., et al., Predicting multiple conformations via sequence clustering and AlphaFold2. Nature, 2024. 625(7996): p. 832-839.

