## Supplementary Information for "Evolutionary Sequence and Structural Basis for the Epistatic Origins of Drug Resistance in HIV"

**Biswas et al.**

**This file includes**

1. **Section S1**: HIV double mutant cycles involving drug-resistance mutations
2. **Table S1**: Frequency of drug-resistance associated positions (DRAPs) among top-ranked DMCs in HIV proteins.
3. **Section S2:** Characterizations of compensatory behavior in literature.
4. **Figure S1**: Key electrostatic interactions contribute to the synergy/antagonism of the mutation pairs.
5. **Section S4**: Structural Basis for the Non-Additive Effects of Highly Synergistic and Antagonistic Integrase Drug Resistance Mutation Pairs
6. **Table S2**: Gain of contacts in top HIV-1 integrase drug resistant double mutants relative to wild-type.
7. Abbreviations List
8. References

### Section S1. HIV double mutant cycles involving drug-resistance mutations

In general, we find that the majority of double mutant cycles (DMCs) within PR and RT involve drug resistance associated positions (DRAPs), whereas ~30-40% of the top 50 cycles within IN involve DRAPs. The simplest explanation for these trends is the relative size of the drug binding site with respect to the whole complex. Both PR and RT are substantially smaller than the viral intasome, and thus coupled mutations are more likely to reside near the ligand binding site and cause drug resistance, thereby becoming DRAPs. Additionally, there may be effects caused by the length of ART therapy. For example, since NRTIs have been around for substantially longer than NNRTIs, this may explain why a greater fraction of DMCs have been sampled during drug resistance evolution. We also note that ~70% of all known DRMs (as defined in Stanford HIVDB) exhibit strong synergistic DMCs. Because NRTIs and NNRTIs distinctly target RT and are both used in ART, we present these data jointly and additionally separate them into the RAPs for the two drug classes. Of note, since the same DRMs can arise under both NRTI and NNRTI treatment, the rows for NRTI/NNRTI DMCs at DRAPs do not necessarily add up to the total DMCs at DRAPs for RT.

**Table S1**: Frequency of drug-resistance associated positions (DRAPs) among top-ranked DMCs in HIV proteins.

| Protein | | % of top 50 DMCs at resistance associated positions | |
|---|---|---|---|
| | | **Synergistic** | **Antagonistic** |
| PR | | 50% | 100% |
| RT | All DRMs | 82% | 68% |
| | NRTI DRMs | 64% | 40% |
| | NNRTI DRMs | 22% | 34% |
| IN | | 40% | 38% |

### Section S2. Characterizations of compensatory behavior in the literature.

In PR the mutation N88D compensates for the reduced replication fitness associated with D30N [1-3], and these are accordingly the strongest coupled DMCs in our analysis. Similarly, I47V is known to act synergistically with V32I to reduce susceptibility to several drugs [1]. We also observe the pairs M46I-

L76V, M46I-V32I, and V32I-L76V. The mutation M46I increases protease catalytic efficiency [3] and occurs in combination with V32I, and L76V [1-3].

In RT, we observe the two distinct patterns (type 1 and type 2) of the thymidine analogue mutations (TAMs) that affect the susceptibility of the NRTI class of drugs by facilitating primer unblocking (aka nucleotide excision) [4, 5] among the top DMCs. For example, we can distinguish between the type 1 and type 2 TAMs, with T215Y-M41L (type 1), and K70R-K219E, D67N-K219Q (type 2) being the top DMCs in concordance with clinical observations [6-8]. We also observe the Q151M pathway complex mutations. Q151M is usually found in combination with two or more of the following accessory mutations: A62V, V75I, F77L, and F116Y [9, 10]. Among mutation pairs selected by the NNRTIs, the pair P225H-K103N are known to occur in combination [11, 12] causing multiple fold resistance to a number of NNRTI drugs [13, 14]. Interestingly, we also discover a new pair (L74V-L100I) amongst the strongest DMCs, where the former is an NRTI-selected DRM, while the latter is an NNRTI-selected resistance mutation.

In IN, the pan-INSTI DRMs arising at G140A/S-Q148H/R/K reduce susceptibility to both 1st and 2nd generation inhibitors [15], and these are accordingly the strongest DMCs in our analyses. Another pair of mutations among the strongest DMCs, T97A-Y143R are known to reduce susceptibility of 1st generation INSTIs by >100 fold in combination [16, 17], a 10 fold increase from Y143R alone. Similarly, E138K/T are annotated to accessory mutations that do not reduce INSTI susceptibility alone [18], but combination with Q148 mutations reduce 10-100−fold [15, 17, 18] and also observed among the strongest DMCs.

**Section S3. Previously unknown antagonistically coupled drug resistance mutation pairs that can be targeted by distinct drugs in combination or sequential order**

We uncover previously unknown highly antagonistically coupled pairs that can be targeted by distinct drugs in combination or sequentially. These involve an NRTI+NNRTI selected pair, as well as pairs of mutations engendering resistance to different PIs and different generation INSTIs. Among the latest generation INSTIs used in ART, Cabotegravir (CAB) acts as a long-acting injectable; however, resistance to CAB occurs through multiple pathways including mutations at Q148. Prior exposure to the 1st generation INSTI Raltegravir (RAL) or presence of N155H mutation in the sequence background can increase resistance barrier to CAB through antagonism with Q148H. Interestingly, the mutation I151V is a highly polymorphic mutation that is found in wild-type molecular clones of HIV such as NL4-3 used in the laboratory which shows antagonism with several DRMs including S147G selected by the 1st generation INSTI, Elvitegravir (EVG), implying that c-ART using EVG can have a higher barrier to resistance for patients with V151I polymorphisms.

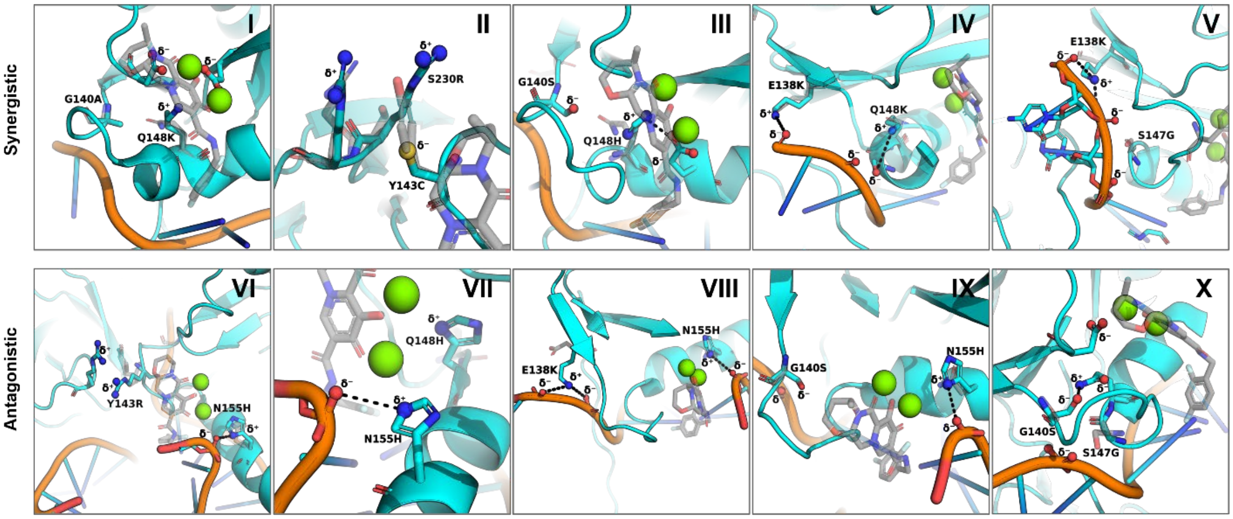


**Figure S1:** The strongest synergistic (I: G140A-Q148K; II: Y143C-S230R; III: G140S-Q148H; IV: E138K-Q148K; V: E138K-S147G) and antagonistic (VI: Y143R-N155H; VII: Q148H-N155H; VIII: E138K-N155H; IX: G140S-N155H; X: G140S-S147G) drug resistant mutation pairings relative to wild-type HIV integrase (gray, PDB: 8FN7). Dolutegravir (gray) and the required $Mg^{+2}$ ions (green) are also shown for reference. Key electrostatic interactions that would contribute to the synergy/antagonism of the mutation pairs are denoted as spheres ($\delta^+$, $\delta^-$). The models used for analysis were taken from experimentally determined structures (8FNL, 8FNM) or from refined models generated (see Methods).

### Section S4: Structural Basis for the Non-Additive Effects of Highly Synergistic and Antagonistic Integrase Drug Resistance Mutation Pairs

Here, we describe in more detail the mechanistic considerations for synergistic coupling between the DMC pairs G140A-Q148H, G140A-Q148K, Y143C-S230R, E138K-Q148K and S147G-E138K.

**G140S–Q148H ($\Delta\Delta E = 8.5$, $\delta = 4.5 \pm 0.5$ kcal/mol)**: Individually, Q148H introduces a positively charged imidazole group at a key catalytic site, near the $Mg^{2+}$ ions coordinated by the Asp64-Asp116-Glu152 motif. This charge perturbs the electrostatic landscape of the active site, reducing the enzyme's catalytic efficiency. On its own, G140S introduces steric bulk and polarity via the hydroxyl-containing serine residue in the flexible catalytic loop, potentially disturbing local geometry, decreasing loop mobility, and introducing repulsive interaction with the DNA phosphate backbone at D17.E. However, when both mutations co-occur, the hydroxyl of Ser140 (bearing a partial negative charge, $\delta^-$) forms an ionic interaction with the positively charged Nδ1 of His148

($\delta^+$), observed within ~4 Å in structural simulations. This interaction effectively neutralizes the charge imbalance caused by Q148H, restoring active site stability and enzyme function. The polar interaction is unique to the double mutant, absent in the respective single mutants. Among all amino acids, Ser is uniquely suited to form this specific interaction. Thr is bulkier and introduces steric hindrance, while Cys has a lower electronegativity and forms weaker hydrogen bonds, which leads to less strong albeit still synergistic couplings with 148H/R/K (ΔΔE ~2-3**)** in comparison to the G140S-Q148H DMC pair. Therefore, this pair represents a structurally optimal and evolutionary preferred combination that maximizes functional compensation, resulting in a strongly synergistic effect. Other frequently observed mutations at 148, like 148R/K, due to their similar chemical nature, also exhibit strong couplings with G140S (**Table 2A**, main text).

**G140A–Q148K (ΔΔE = 5.6, δ = 3.5 ± 0.78 kcal/mol)**: Q148K, when present alone, introduces a large, flexible, positively charged lysine side chain into a constrained region near the catalytic core, which disrupts loop conformation and also changes the local environment near the catalytic domain, leading to the reorientation of His114 from its stable initial position and reducing the enzymatic activity[19]. G140A, in contrast, introduces a bulkier group at position 140 by replacing glycine with alanine. On its own, G140A increases loop rigidity and alters side chain packing, which is somewhat detrimental[19]. In the double mutant, the presence of alanine at position 140 allows Lys148 to rotate +115° around its γ-carbon, repositioning the positively charged side chain closer to the negatively charged residues of the DDE triad. This repositioning partially mitigates a new electrostatic interaction with the $Mg^{2+}$-coordinating DDE residues, thus restoring enzymatic function to a limited degree that is severely disrupted by Q148K alone. Although the enzymatic activity and viral replicative capacity of the double mutant is still compromised, the double mutant exhibits compensatory epistasis that mitigate the detrimental effects anticipated from the additive combination of the two single mutations. Structurally, other substitutions at position 140 (e.g., bulkier polar residues) would restrict this conformational flexibility, making G140A the minimal and necessary substitution for this compensatory mechanism. This precise geometric accommodation underlies the observed synergistic interaction. A similar interaction is also observed with the Q148R and Q148H mutations, both of which exhibit strong synergistic epistatic coupling with G140A. (**Table 2A**).

**Y143C–S230R (ΔΔE = 6.5, δ = 3.93 ± 0.98 kcal/mol)**: In the wild-type IN, residues Y143 and R2301 may engage in an inter protomer π-cation interaction between the aromatic ring of Tyr143 and the -$NH_3^+$ of Arg, contributing to structural stability. The Y143C mutation replaces the bulky, rigid tyrosine with a more flexible cysteine, thereby disrupting this interaction and introducing a potentially repulsive environment between Cys143 and Ser230, which could weaken the packing that is necessary for intasome formation. On the other hand, the S230R mutation introduces a large, positively charged arginine at the C-terminal region. This substitution alone may create steric clashes with Tyr143, destabilizing the inter-subunit interface. Remarkably, when these mutations co-occur, they give rise to a unique structural adaptation: the thiol (-SH) group of Cys143 should form an ionic interaction (~4 Å) with the guanidinium (-$NH_3^+$) group of Arg230 across adjacent protomers. This inter-chain interaction would compensate for the destabilizing effects of the individual mutations alone and reinforces the inter-protomer interface within the intasome complex. Notably, this stabilizing interaction is only feasible due to the specific chemical properties of cysteine or serine at position 143 and arginine at position 230. Other commonly observed substitutions at position 143, such as His or Lys (143H/K), would likely repel Arg230 due to charge similarity, while Ala or Gly (143A/G) would lack sufficient polarity to form productive interactions. Although serine at 143 shares a similar size with cysteine, its higher electronegativity may lead to an overly strong interaction with Arg230, potentially leading to weaker synergistic couplings (ΔΔE = 3.70). Due to the size and charge similarity between Arg and Lys, S230K exhibit similar but less stronger interactions with Y143C, which leads to stronger couplings (ΔΔE = 3.03). Thus, the Y143C–S230R pair represents a uniquely strong synergistic interaction enabled by a compensatory cross-interface salt bridge that is not as strong with alternative residue combinations.

**E138K–Q148K (ΔΔE = 5.0, δ = -3.29 ± 0.80 kcal/mol)**: When present alone, the Q148K mutation introduces a large, flexible, and positively charged lysine side chain into a spatially constrained region near the catalytic site of the IN enzyme. This not only disrupts the conformation of the catalytic loop but also alters the local electrostatic environment, impairing loop flexibility and misaligning the critical His114 residue that is essential for catalysis. E138K alone reorients to form a salt-bridge contact with the backbone phosphate of DNA, which disrupts the internal loop packing. When combined, E138K compensates for Q148K-induced rigidity by restoring loop flexibility and aligning His114 appropriately. This is well-known from high-resolution structural data[19]. Despite the lack of direct interactions between the two mutated residues, their opposing effects on loop dynamics are complementary. Functionally, E138K allows Q148K to maintain catalytic activity while avoiding the detrimental conformational freeze. This presumed balance of rigidity and flexibility highlights the cooperative nature of this pair, leading to a structurally and energetically synergistic outcome. Other commonly observed mutations at position 148, such as Q148R/H, also exhibit strong coupling with E138K, likely due to the similar chemical properties of arginine, histidine, and lysine. In contrast, substitutions at position 138 such as E138A or E138T do not show strong coupling with Q148R/K/H, as these residues lack the ability to form salt bridge interactions with the DNA backbone. Consequently, they are unable to compensate for the loop rigidity induced by Q148R/K/H or to restore the proper orientation of His114, which is critical for maintaining catalytic function.

**E138K–S147G ($\Delta\Delta E$ = 2.6, $\delta$ = -1.67 ± 0.71 kcal/mol)**: E138K, on its own, disturbs internal loop packing by establishing a salt-bridge interaction with the DNA phosphate backbone at nucleotide D16.E. S147G removes the polar hydroxyl group of Ser147, which normally participates in stabilizing electrostatic interactions with DNA. This change increases the flexibility of the catalytic loop, facilitating loop breathing and allowing E138K to adopt a favorable conformation for its DNA interaction. When combined, these mutations improve DNA binding affinity through enhanced adaptability of the loop. The structural synergy stems from the complementary nature of E138K's stabilizing function and S147G's dynamic enhancement, neither of which is fully realized in the single mutants. The bulkier Ala147 may impart greater rigidity towards the loop in comparison to Gly147. Other mutations such as 138A/T, will not form productive salt bridge interactions with the DNA phosphate, which Gly147 can facilitate.

Here, we describe in more detail the mechanistic considerations for antagonistic coupling between the DMC pairs Y143R+N155H, E138K-N155H, Q148H-N155H, G140S-N155H and S147G-G140S.

**Y143R–N155H ($\Delta\Delta E$ = -2.31, $\delta$ = 3.06 ± 0.64 kcal/mol)**: Alone N155H forms a salt bridge with the nearby DNA phosphate group at position D21.F. It may additionally critically disrupt the catalytic site by intruding upon and forming a new salt bridge with Glu152, destabilizing the enzyme active site. Meanwhile, Y143R itself introduces a positively charged arginine that causes electrostatic repulsion with Arg231 on the opposing protomer. Together, the detrimental effect on the active site, including mis-coordinated DNA contacts, as well as the repulsion between Arg143 and Arg231, would likely generate a distorted electrostatic field and structural instability. These effects amplify each other's detrimental influence on protein fitness, resulting in a strongly antagonistic interaction that severely disrupts catalytic efficiency. Y143K/H share similar chemical properties with Y143R but have slightly lower positive charge, leading to less disruptive effects when combined with N155H. However, both the Y143K/H–N155H DMC pairs remain antagonistic (ΔΔE = -1.65).

**Q148H–N155H (ΔΔE = -2.31, δ = 3.47 ± 0.53 kcal/mol)**: Individually, Q148H introduced a bulky positively charged group near the catalytic site, which alters the electrostatic landscape of the active site, reducing the enzyme's catalytic efficiency. As described above, the effect of N155H may be dually disruptive due to the formation of a new interaction with the DNA phosphate on D21.F and its intrusion into the active site via interaction with Glu152. Together, each mutation creates opposing interactions that give rise to structural tension across the active site, akin to a molecular tug-of-war. The structural polarity imposed by these mutations causes distortion of the catalytic geometry, reducing the ability of integrase to effectively coordinate DNA and $Mg^{2+}$ ions. Because the mutations target opposing arms of the active site, their combined effect creates destabilizing vector forces that interfere with one another. The result is a non-additive, antagonistic outcome that is further destabilizing in comparison to the additive effects of each individual mutation alone. Q148R/K exhibits stronger antagonistic coupling with N155H (**Table 2B**, main text), which may be attributed to the similar chemical properties shared among the H, R, and K residues. Limited space near the active site likely prevents accommodation of bulkier positively charged amino acid (Lys/Arg) at position 155.

**E138K–N155H (ΔΔE = -1.2, δ = 1.7 ± 0.89 kcal/mol)**: E138K disturbs internal loop packing by introducing a salt-bridge interaction with DNA. As above, N155H is dually disruptive. The two mutations act on the same viral DNA strand, separated by five nucleotides in sequence. The overlapping regions of influence between these mutations likely lead to a misaligned loop and opposing DNA strain. Together, these mutations would misalign active site flexibility via destabilization of DNA contacts that are necessary for catalysis and interfering with optimal catalytic loop flexibility. These conflicting structural demands lead to antagonism, and any potential compensatory benefits from E138K alone, which is occasionally observed in viral infectivity assays[19], is counteracted by the adverse effects of one another.

**G140S–N155H (ΔΔE = -2.12, δ = 2.19 ± 0.77 kcal/mol)**: G140S adds steric bulk and polarity through the hydroxyl group of the serine residue within the flexible catalytic loop, potentially disrupting local geometry and reducing loop mobility. Additionally, it creates a repulsive interaction with the DNA phosphate backbone at D17.E, while N155H introduces an attractive interaction with DNA at D21.F. These opposing interactions occur on different DNA strands, producing diverging vector forces that increase DNA tension and misalignment within the intasome complex. The resulting structural stress would disrupt the DNA–protein interface and weaken their interactions. While each mutation individually maintains partial DNA engagement, their combination leads to geometric distortion of the substrate vDNA, rendering the pair functionally incompatible and antagonistic. Cys, being less electronegative results in weaker couplings, although the G140C-N155H DMC pair notably remains antagonistic (ΔΔE = -1.02).

**G140S–S147G (ΔΔE = -2.10, δ = 1.37 ± 0.47 kcal/mol):** Both G140S and S147G independently disrupt DNA binding at D17.E. G140S introduces a negatively charged hydroxyl group that adds steric bulk and polarity withing the catalytic loop and also repels the DNA phosphate, while S147G removes a stabilizing polar contact from the loop. The combined effect is a dual destabilization of the same DNA contact site, resulting in weakened DNA interaction and disordered catalytic loop architecture. This redundancy in functional disruption causes overlapping negative effects that outweigh any compensatory benefits, leading to structural and functional antagonism within this DMC pair. Both Cys (ΔΔE = -1.95) and Thr (ΔΔE = -1.83) at position 140, due to their similar chemical nature but weaker electronegativity (Cys) and slightly bulkier side chains (Thr), exhibit weaker albeit still antagonistic behavior when coupled with S147G.

These analyses highlight the structural mechanisms underlying both synergistic and antagonistic effects of integrase drug resistance mutations, shedding light on their impact on the intrinsic protein stability. Similar considerations may apply more generally to other proteins.

**Table S2**: Gain of contacts in top HIV-1 integrase drug resistant double mutants relative to wild-type.

This table summarizes the changes in molecular contacts, specifically DNA, inter-protein, and intra-protein interactions, upon transitioning from wild-type (WT) to selected synergistic or antagonistic double mutant pairs. Contact differences ($\Delta$WT→M1M2) represent the net gain in each category compared to WT, based on structural analysis. Synergistic mutation pairs often show increased intra-protein stabilization or restored DNA interactions, while antagonistic pairs also exhibit contact gains, but may involve conflicting structural adjustments that contribute to non-additive or disruptive outcomes. The contacts are calculated using experimental and modelled structures (See methods)

| **Mutation Pair** | $\Delta_{WT \rightarrow M1M2}$ **DNA contacts** | $\Delta_{WT \rightarrow M1M2}$ **Inter-protein contacts** | $\Delta_{WT \rightarrow M1M2}$ **Intra-protein contacts** |
|---|---|---|---|
| **Synergistic Pairs** | | | |
| E138K/Q148K | +1 | 0 | +1 |
| G140A/Q148K | 0 | 0 | +2 |
| G140S/Q148H | 0 | 0 | +2 |
| Y143C/S230R | 0 | +1 | 0 |
| E138K/S147G | +2 | 0 | 0 |
| **Antagonistic Pairs** | | | |
| Y143R/N155H | +1 | 0 | 0 |
| Q148H/N155H | +1 | 0 | +2 |
| G140S/N155H | +1 | 0 | +2 |
| G140S/S147G | +1 | 0 | +1 |
| E138K/N155H | +2 | 0 | +1 |

**List of Abbreviations Used**

| Abbreviation | Full Form |
|---|---|
| 3TC | Lamivudine |
| ABC | Abacavir |
| ART | Antiretroviral Therapy |
| ARV | Antiretroviral |
| ATV | Atazanavir |
| AZT | Zidovudine (also known as Retrovir) |
| BAR | Bennett’s Acceptance Ratio |
| CAB | Cabotegravir |
| Ca | Calcium |
| DMC | Double Mutant Cycle |
| DRM | Drug Resistance Mutation |
| DRV | Darunavir |
| EFV | Efavirenz |
| ETR | Etravirine |
| EVG | Elvitegravir |
| FEP | Free Energy Perturbation |
| FTC | Emtricitabine |
| HIV | Human Immunodeficiency Virus |
| HIVDB | HIV Drug Resistance Database |
| IN | Integrase |
| INSTI | Integrase Strand Transfer Inhibitor |
| LPV | Lopinavir |
| MD | Molecular Dynamics |
| MSA | Multiple Sequence Alignment |
| NFV | Nelfinavir |
| NNRTI | Non-Nucleoside Reverse Transcriptase Inhibitor |

| | |
|---|---|
| NRTI | Nucleoside Reverse Transcriptase Inhibitor |
| NVP | Nevirapine |
| PI | Protease Inhibitor |
| PR | Protease |
| RAL | Raltegravir |
| RPV | Rilpivirine |
| RT | Reverse Transcriptase |
| SD | Single Domain |
| TAMs | Thymidine Analogue Mutations |
| TDF | Tenofovir Disoproxil Fumarate |
| TFV | Tenofovir |
| TPV | Tipranavir |
| WT | Wild Type |
| c-ART | Combination Antiretroviral Therapy |
| vDNA | Viral DNA |
| λ-REMD | Lambda-Replica Exchange Molecular Dynamics |

**References:**


1. Rhee, S.-Y., et al., *HIV-1 protease mutations and protease inhibitor cross-resistance.* Antimicrobial agents and chemotherapy, 2010. **54**(10): p. 4253-4261.
2. Vermeiren, H., et al., *Prediction of HIV-1 drug susceptibility phenotype from the viral genotype using linear regression modeling.* Journal of virological methods, 2007. **145**(1): p. 47-55.
3. Henderson, G.J., et al., *Interplay between single resistance-associated mutations in the HIV-1 protease and viral infectivity, protease activity, and inhibitor sensitivity.* Antimicrobial agents and chemotherapy, 2012. **56**(2): p. 623-633.
4. Boyer, P.L., et al., *Selective excision of AZTMP by drug-resistant human immunodeficiency virus reverse transcriptase.* Journal of virology, 2001. **75**(10): p. 4832-4842.
5. Sluis-Cremer, N., D. Arion, and M. Parniak*, *Molecular mechanisms of HIV-1 resistance to nucleoside reverse transcriptase inhibitors (NRTIs).* Cellular and Molecular Life Sciences CMLS, 2000. **57**: p. 1408-1422.

6. Miller, M.D., et al., *Genotypic and phenotypic predictors of the magnitude of response to tenofovir disoproxil fumarate treatment in antiretroviral-experienced patients.* Journal of Infectious Diseases, 2004. **189**(5): p. 837-846.
7. De Luca, A., et al., *Frequency and Treatment-Related Predictors of Thymidine-Analogue Mutation Patterns in HIV-1 Isolates after Unsuccessful Antiretroviral Therapy.* The Journal of Infectious Diseases, 2006. **193**(9): p. 1219-1222.
8. Rhee, S.-Y., et al., *Distribution of Human Immunodeficiency Virus Type 1 Protease and Reverse Transcriptase Mutation Patterns in 4,183 Persons Undergoing Genotypic Resistance Testing.* Antimicrobial Agents and Chemotherapy, 2004. **48**(8): p. 3122-3126.
9. Shafer, R.W., et al., *Drug Resistance and Heterogeneous Long-Term Virologic Responses of Human Immunodeficiency Virus Type 1-Infected Subjects to Zidovudine and Didanosine Combination Therapy.* The Journal of Infectious Diseases, 1995. **172**(1): p. 70-78.
10. Iversen, A.K., et al., *Multidrug-resistant human immunodeficiency virus type 1 strains resulting from combination antiretroviral therapy.* Journal of Virology, 1996. **70**(2): p. 1086-1090.
11. Bacheler, L.T., et al., *Human Immunodeficiency Virus Type 1 Mutations Selected in Patients Failing Efavirenz Combination Therapy.* Antimicrobial Agents and Chemotherapy, 2000. **44**(9): p. 2475-2484.
12. Alcaro, S., et al., *Docking Analysis and Resistance Evaluation of Clinically Relevant Mutations Associated with the HIV-1 Non-nucleoside Reverse Transcriptase Inhibitors Nevirapine, Efavirenz and Etravirine.* ChemMedChem, 2011. **6**(12): p. 2203-2213.
13. Bacheler, L., et al., *Genotypic Correlates of Phenotypic Resistance to Efavirenz in Virus Isolates from Patients Failing Nonnucleoside Reverse Transcriptase Inhibitor Therapy.* Journal of Virology, 2001. **75**(11): p. 4999-5008.
14. Smith, S.J., et al., *Rilpivirine and Doravirine Have Complementary Efficacies Against NNRTI-Resistant HIV-1 Mutants.* JAIDS Journal of Acquired Immune Deficiency Syndromes, 2016. **72**(5): p. 485-491.
15. Cahn, P., et al., *Dolutegravir versus raltegravir in antiretroviral-experienced, integrase-inhibitor-naive adults with HIV: week 48 results from the randomised, double-blind, non-inferiority SAILING study.* The Lancet, 2013. **382**(9893): p. 700-708.
16. Blanco, J.-L., et al., *HIV-1 integrase inhibitor resistance and its clinical implications.* Journal of Infectious Diseases, 2011. **203**(9): p. 1204-1214.
17. Canducci, F., et al., *Cross-resistance profile of the novel integrase inhibitor Dolutegravir (S/GSK1349572) using clonal viral variants selected in patients failing raltegravir.* Journal of Infectious Diseases, 2011. **204**(11): p. 1811-1815.
18. Kobayashi, M., et al., *In vitro antiretroviral properties of S/GSK1349572, a next-generation HIV integrase inhibitor.* Antimicrobial agents and chemotherapy, 2011. **55**(2): p. 813-821.
19. Li, M., et al., *Mechanisms of HIV-1 integrase resistance to dolutegravir and potent inhibition of drug-resistant variants.* Sci Adv, 2023. **9**(29): p. eadg5953.
